# Speed-Dependent Turning Strategies in Quadrupedal Locomotion: Insights from Computational Modeling

**DOI:** 10.64898/2026.01.12.699101

**Authors:** Yaroslav I. Molkov, Mohammed A. Y. Mohammed, Tommy Stell, Amelia Harralson, Russell Jeter, Ilya A. Rybak

## Abstract

Quadrupedal animals like mice navigate their environments through complex coordination of neural signals and biomechanical movements, enabling stable and directed locomotion. While many computational models simplify this process by assuming left-right symmetrical body movements and focusing on straight-line paths, real animals rely heavily on asymmetrical body movements to execute turns and adjust speed effectively. This study builds upon a previously developed model of quadrupedal locomotion proposed by Molkov et al., 2024) in which forward movement of the body was driven by central neural interactions, biomechanics, and proprioceptive feedback. We extended this model to comparatively investigate possible mechanisms of steering by introducing three distinct asymmetrical strategies–body bending, lateral force application, and lateral limb shifting as well as their combinations–to explore their potential involvement in turning performance. By simulating these strategies across a walking speed range, we measured and compared their impact on turning curvature the sharpness of the turn) and limb coordination. The latter was quantified through ratios of duty factors representing the relative time that a limb spent in contact with the ground compared to its counterpart on the opposite side. Our findings reveal that each strategy excels at different speeds: body bending allows sharp turns at low speeds, lateral force is most effective at medium speeds, and lateral shifting performs best at higher speeds. Our results suggest that animals select or combine turning strategies based on their locomotor speed or adjust speed to use a specific strategy. We also show that the forelimbs consistently play a primary role in steering, while the hindlimbs adjust propulsion and stability in ways that depend on the specific turning strategy. These results provide valuable insights into how spinal circuits and mechanical asymmetries work together to produce flexible, adaptive movement patterns, offering a robust framework for understanding locomotion in both biological organisms and robotic systems designed to mimic such behaviors.

## Introduction

Quadrupedal locomotion is characterized by a remarkable coordination of speed and limb movements, allowing animals like mice to adaptively change direction and navigate in complex and varied environments. This ability relies on the interplay between the central nervous system, which includes Central Pattern Generators (CPGs) generating rhythmic motor patterns (for review, see Grillner, 2006; Orlovskiĭ, 1999; McCrea et al., 2008; Stuart et al., 2008; Rybak et al., 2015), and the musculoskeletal system that translates these motor patterns into physical motion (Orlovskiĭ, 1999; Bertram, 2016). While straight-line locomotion has been extensively studied, the mechanisms governing turning–a critical maneuver that requires breaking bilateral symmetry–are less well understood. Many computational models of quadrupedal locomotion, including those based on CPGs (e.g. Danner et al., 2016; Danner et al., 2017; Rybak et al., 2024; Rybak et al., 2025), assume a symmetrical body and focus on forward movement. However, these simplifications fail to capture the dynamic asymmetries that real animals employ during turning, such as adjusting body posture or limb forces to change direction (Pycock, 1980;, Walter, 2003; Gruntman et al., 2007; Cregg et al., 2020; Usseglio et al., 2020; Haagensen et al., 2022).

Turning is essential for survival, enabling animals to avoid obstacles, pursue prey, or evade predators. Unlike straight-line movement, turning demands the generation of asymmetrical forces and movements, which arise from intricate left-right asymmetric interactions between neural commands and biomechanical structures (Cregg et al., 2020; Usseglio et al., 2020). For instance, mice may bend their spines or reposition their limbs to execute sharp turns, strategies that are actively controlled rather than mere deviations from symmetry (Walter, 2003; Gruntman et al., 2007). Understanding these asymmetrical strategies is crucial for developing computational models that more accurately reflect biological locomotion and for informing the design of agile robotic systems capable of navigating complex terrains (Wei et al., 2018; Lee et al., 2020).

Here we focus on mouse-like turning during walking in a relatively low-speed regime, where support geometry, limb loading, and stride timing can be analyzed within a planar approximation. Faster gaits with aerial phases, pronounced roll or pitch, and broader mass-dependent three-dimensional effects are outside the scope of the present model. To investigate turning behaviors, we extended a simplified model of quadrupedal locomotion, originally developed by (Molkov et al., 2024). The original model simulated straight-line walking in mice using sensory feedback to the CPG-based central neural controller to generate limb movements. Turning in this framework is produced by controlled biomechanical asymmetries. We examine three minimal and mechanistically distinct routes to directional change: body bending, explicit lateral force generation, and asymmetrical foot placement. These strategies were selected not because they exhaust the turning repertoire of animals, but because each isolates a different way of breaking left-right symmetry in a simplified neuromechanical framework. This design allows direct comparison of how geometric, dynamic, and kinematic asymmetries shape turning curvature and limb coordination as speed changes. Specifically, we measured curvature, which indicates the sharpness of a turn and characterized the dependence of the maximal achievable curvature on speed and the choice of a turning strategy. To investigate the related changes in limb coordination during turning, we analyzed duty factor ratios, defined as the ratio of the proportion of time a limb spends in contact with the ground (stance phase) relative to its total stride cycle for the left versus right limbs. Our objectives were to determine how these basic turning strategies influence turning performance at different speeds and to explore the distinct roles of forelimbs and hindlimbs in achieving directional control. By doing so, we aim to elucidate the interplay between spinal circuits and mechanical asymmetries, providing insights applicable to both biological locomotion and robotic system design. The findings offer insights into the neuromechanical mechanisms that quadrupedal animals employ for steering and navigation while moving.

## Model Description

The computational model used in this study is grounded in biomechanical principles observed in animal locomotion, emphasizing the role of proprioceptive feedback in control of locomotion (Pfeifer et al., 2007; Frigon et al., 2021; Molkov et al., 2024; Rybak et al., 2024). Unlike models that rely on pre-programmed neural oscillators, this model posits that complex rhythmic movements emerge from distributed sensory feedback loops and the intrinsic mechanical properties of the body (Full et al., 1999; Holmes et al., 2006); a concept also known as “morphological computation” (Owaki et al., 2013; Müller et al., 2017). The model is intentionally reduced to three planar degrees of freedom (x, y, and yaw) to isolate the core mechanics of turning while keeping the system analytically interpretable and computationally tractable. This simplification excludes roll, pitch, aerial phases, and elastic trunk deformation, and therefore the model should be interpreted as a minimal representation of mouse-like walking rather than a full biomechanical reconstruction of quadrupedal turning.

### Model of the body

In our model, the mouse body is represented as a rigid rod of length *L* possessing uniform linear density, oriented parallel to the ground at distance *H* (height of the mouse), and having three degrees of freedom: horizontal position (x, y coordinates) and orientation (Fig. 1). In the horizontal plane, the left and right “shoulder” and “hip” joints are equidistant, at a distance *h* from the rod, forming the body frame, a *L* x 2*h* area rectangle. The center of mass (COM) is located in the middle of the body.

**Figure 1:**
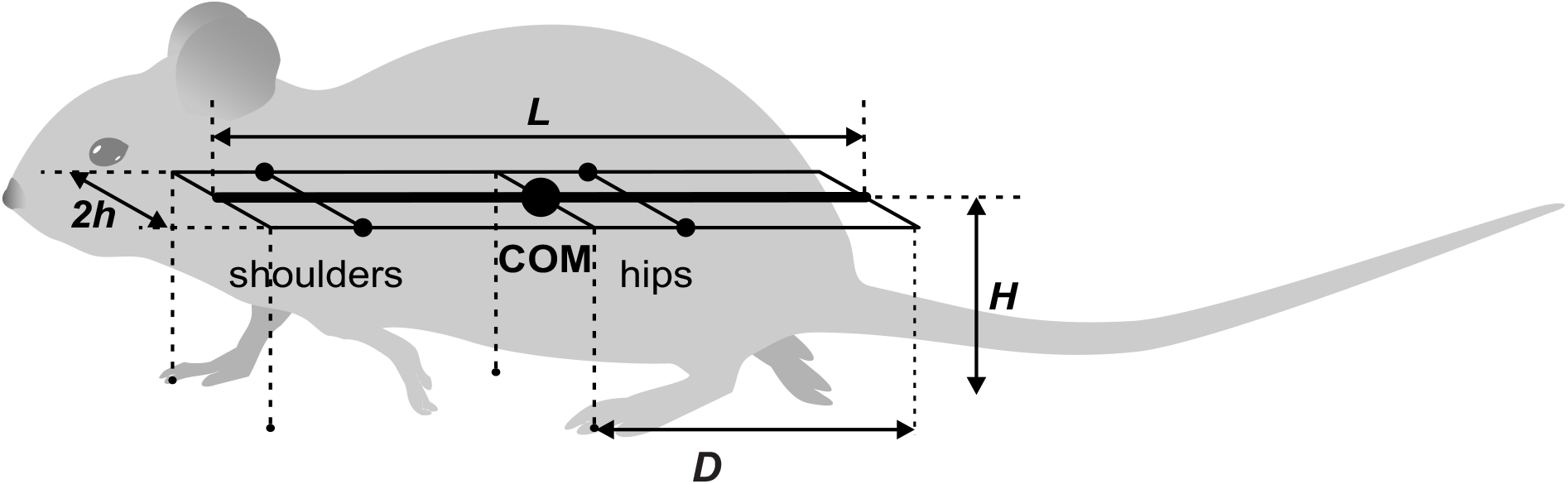
The mouse’s body as represented in the model

The body’s weight is supported by its limbs during stance. The initial paw positions are defined by the ground projections of the corners of the body frame as indicated by the vertical dashed lines in Fig. 1: the front of the frame for the forelimbs and the center (aligned with the COM) for the hindlimbs (Fig. 1). These positions, which are laterally displaced by *h* from the centerline, also serve as the default target positions for the corresponding paws during the swing phase. Here *D* = *L*/2 denotes the maximal two-dimensional displacement magnitude of a paw from its touchdown reference position in body co-ordinates, rather than only the axial component of that displacement. Consequently, a mediolateral off-set can cause the displacement limit to be reached earlier even when the axial component of stance excursion is reduced.

### Central controller and control of movement

The model has a central controller or Central Pattern Generator (CPG) consisting of four Rhythm Generators (RGs) each controlling one limb and operating as a state machine with two states defining the stance (ground contact) and swing (off-ground movement) phases of the corresponding limb. Sensory feedback signals that characterize limb loading, limb extension, and overall postural stability modulate these generators (by controlling the timing of stance-to-swing and swing-to-stance transitions) enabling adaptive locomotion (for details see Methods and (Molkov et al., 2024)).

Movement in the model arises from the interaction between a central neural controller and the mechanical components of the system. Each limb alternates between stance, where it contacts the ground to generate propulsion, and swing, where it moves freely to prepare for the next step. Load-sensitive receptors detect when a limb is bearing weight or unloading, directly influencing the timing of phase transitions and ensuring balance and forward motion.

During the swing phase, limbs do not contribute to propulsion. Limb loading is the vertical component of the ground reaction force which is calculated from the COM coordinates relative to limb positions. A limb transitions from the stance to swing phase when its loading becomes zero or negative, or its extension limit is surpassed. Swing-to-stance transitions are governed by a feedback control mechanism to prevent loss of balance (see (Molkov et al., 2024) and Methods).

The model’s equations of motion describe the dynamics of the mouse body in a horizontal plane, treating it as a rigid system with a center of mass (COM). These equations are derived from Newton’s second law applied to the planar motion of the COM and to the angular momentum for the body’s rotation around the COM. They incorporate kinematic friction to account for energy dissipation during locomotion. The primary forces acting on the system include propulsion forces generated by the limbs in contact with the ground and an inverted pendulum force that arises specifically during phases when only two limbs are supporting the body.

Ground reaction forces, which represent the interaction between the paws and the ground, are decomposed into **horizontal** and **vertical** components. The horizontal components drive forward movement and rotation, while the vertical components support the body’s weight.

## Horizontal forces and yaw torques

In the horizontal plane, each limb in stance generates a propulsion force of equal magnitude for all supporting limbs, directed along the paw’s displacement from its initial position during stance in the body’s coordinate system. This force acts as a key control parameter that influences the overall locomotor speed. To model energy losses from various unrepresented factors like muscle inefficiencies or ground interactions, a viscous friction force is included, which opposes the COM’s velocity and scales linearly with it.

When only two limbs remain in stance, the body cannot in general satisfy the full static support conditions assumed for three- or four-limb support. In this regime we use a planar effective-force approximation: the body is treated as an inverted pendulum about the support line connecting the two grounded paws, and the deviation of the center of mass from that line is represented as a horizontal effective force acting perpendicular to the support line. We refer to this term as the “inverted-pendulum” force to emphasize that it is a reduced planar representation of the destabilizing effect of gravity rather than a direct three-dimensional contact force. During stance, grounded paws remain fixed in ground coordinates until liftoff, while the body moves relative to those fixed contacts. For rotational dynamics, propulsion forces create yaw torques because they are offset from the body’s COM.

### Vertical forces

The vertical components of the ground reaction forces, referred to as “limb loads” or “weight-bearing forces”, support the body’s weight and are computed based on the number of limbs in stance.

For support by more than two limbs, the total vertical force must equal the gravitational force on the body to prevent vertical movement. In the multilimb support calculations, this requirement follows directly from the assumption that body height *H* is fixed and the body has no vertical acceleration. Additionally, the torques from these forces about the COM must balance to avoid pitch or roll, consistent with the assumption that the body frame remains in a horizontal plane. This setup forms a system of balance conditions: force equilibrium and two torque equilibrium equations (for the two horizontal directions). For three-limb support, this yields a unique distribution of loads. Under four-limb support the balance equations are underdetermined, so we select the solution that distributes load as evenly as possible. This additional criterion is a regularizing assumption used to obtain a unique solution; it should not be interpreted as a claim that biological animals necessarily equalize load in the same way. Loads must be positive for limbs to remain on the ground; if a load becomes zero or negative, it indicates unloading and triggers a transition to swing. Geometrically, positive loads require the COM to lie inside the support polygon formed by the paws which is a prerequisite of static stability.

For three- and four-limb support, vertical loads are computed from force and torque balance under the assumption that body height H remains constant and the body frame remains horizontal. During two-limb support, however, those full static balance conditions are no longer imposed within the planar approximation, so total vertical load need not remain equal to body weight and can decrease as the COM moves away from the equilibrium configuration relative to the support line. We therefore use the onset of decreasing total vertical load as an operational signal that the current support state is becoming unsustainable and that a swing-to-stance transition should be triggered. The corresponding derivation and notation are given in Methods, under Weight bearing forces (Eqs. (3)–(6); Fig. 12A) and Swing-to-stance transition.

For full mathematical details, including derivations and parameter justifications, see Methods. In its baseline configuration, without intentional asymmetries, the model produces stable, symmetrical gaits that result in straight-line locomotion, consistent with typical walking patterns in mice (Molkov et al., 2024).

### Asymmetries for turning

To investigate turning, we extended this model by introducing three distinct forms of controlled asymmetry, designed to mimic strategies observed in real quadrupeds (Fig. 2). First, **Body Bending** allows the hips and shoulders to deviate from parallel alignment, simulating spinal flexion that reorients the body’s axis. Second, **Lateral Force** applies forces perpendicular to the body axis, primarily through the forelimbs, to induce rotational moments. Third, **Lateral Shift** offsets the side-to-side placement of limbs relative to the body’s midline, altering the distribution of ground reaction forces. These modifications enable the model to break from symmetrical gaits and generate turning behaviors, providing a platform to study how asymmetries influence locomotor control.

**Figure 2:**
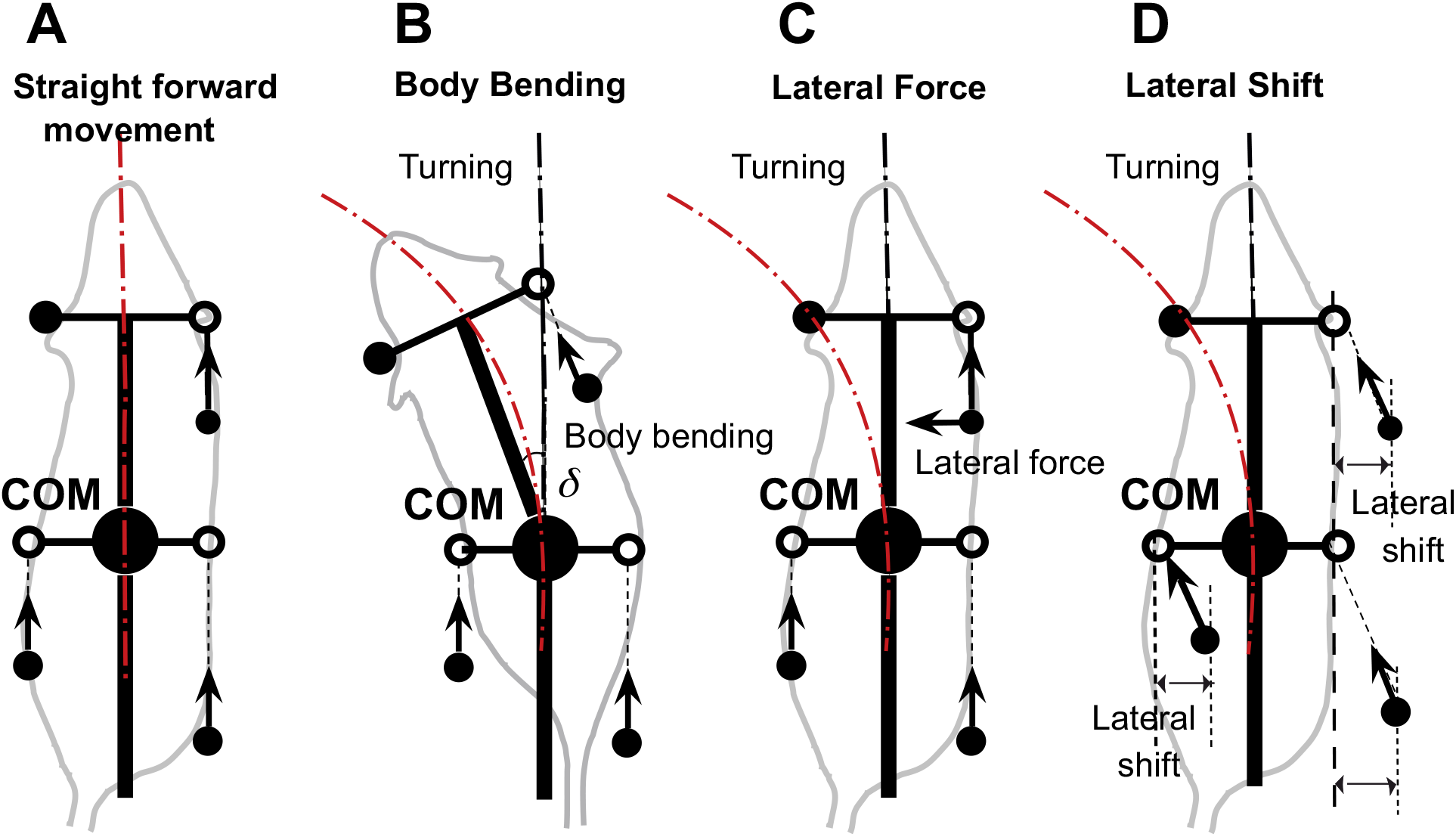
Straightforward movement (**A**) and three turning strategies based on body bending (**B**), lateral force (**C**), and lateral shift (**D**). In the baseline configuration, the model produces a stable, symmetrical gait resulting in straight-line locomotion (**A**). Body bending introduces asymmetry through the non-parallel alignment of the hip and shoulder axes (simulating axial flexion), which shifts the center of mass inward and redistributes propulsive forces to facilitate turning (**B**). The lateral force strategy involves applying a force perpendicular to the body axis, specifically at the forelimbs, to directly generate centripetal acceleration and active steering (**C**). Lateral shift is achieved by displacing the limb landing positions side-to-side relative to the body axis during the swing phase (**D**) This alters the base of support and effective lever arms for ground reaction forces, inducing lateral acceleration and rotation. Black filled circles represent the positions of limbs that are currently in stance phase (ground contact). Open circles indicate default target positions for the limbs during the swing phase. Arrows indicate the directions of propulsive forces.

To explore turning behaviors, we systematically applied each asymmetrical strategy–body bending (Fig. 2B), lateral force (Fig. 2C), and lateral shift (Fig. 2D) –to the computational model. This model extends the original proprioceptive feedback-driven framework for quadrupedal locomotion described in (Molkov et al., 2024). In the original model, the propulsion force generated by each limb during stance was directed strictly along the body’s longitudinal axis, assuming symmetrical and forward-oriented paw displacements. In this extended model, however, the propulsion force is instead directed along the displacement of the paw relative to its unperturbed initial position during stance in the body coordinate system.

This modification allows for more flexible force vectors that can incorporate lateral components in asymmetric conditions, better capturing turning dynamics. In the extended model, the open (white) circle in Fig. 2 denotes the nominal body-fixed target position inherited from the baseline straight-walking model. These target positions reflect the nominal limb-frame geometry relative to the trunk. During stance, propulsion is assumed to arise from active limb extension, so the resulting force acts opposite to the extension vector, equivalently along paw displacement relative to the nominal target. Consequently, a mediolateral shift of touchdown produces a lateral force component opposite to the direction of the shift. For this reason, generating an inward steering force requires shifting paw placement opposite to the intended turn. All propulsion forces remain equal in magnitude across limbs but vary in direction based on actual paw movement, which emerges from the interplay of neural feedback and mechanical constraints. Because the trunk is modeled as a rigid planar body, fore-hind differences in axial force components do not produce explicit trunk compression or stretch in the model; instead, those forces contribute only to the net rigid-body translation and rotation. Under lateral shift, lateral propulsion components arise because the model explicitly defines propulsion direction by paw displacement relative to the nominal target; this differs from the separate Lateral Force strategy, in which an additional perpendicular force component is imposed directly.

Specifically, the asymmetries were introduced as follows:

#### Body bending (Fig. 2B)

This strategy simulates spinal flexion by introducing an angular deviation *δ* between the front (shoulder) and rear (hip) segments of the body. In the base model, the body is a single rigid rod with fixed parallel alignment of hips and shoulders. This alters the relative orientations of the limb attachment points: forelimb positions (ground projections at the front of the frame) and hindlimb positions (at the center) are recalculated in the body’s coordinate system, shifting the target paw positions during swing and the support polygon during stance. Consequently, this reorients the overall body axis over time, generating a rotational torque through asymmetric ground reaction forces without changing force magnitudes. The modified paw displacements under body bending influence the propulsion force directions, potentially introducing slight lateral components.

#### Lateral force (Fig. 2C)

This involves adding a perpendicular component to the propulsion forces, which in the base model are determined by paw displacements but effectively axial in symmetric gaits. The lateral force was applied perpendicular to the body axis, directed inward toward the turn (e.g., leftward for a left turn). This was targeted at the forelimbs to mimic steering. Hindlimbs received zero lateral component to emphasize forelimb-driven rotation. This addition affects the horizontal equations of motion by introducing asymmetric yaw torques (computed as the cross product of the force and the moment arm from the COM to the paw position), leading to rotational acceleration while preserving total forward propulsion.

#### Lateral shift (Fig. 2D)

This kinematic asymmetry adjusts the side-to-side limb placements relative to the body’s midline. In the base model, left and right limbs are symmetrically displaced by *h* (half-width) from the centerline. To implement the shift, an off-set *s* was applied asymmetrically: for a left turn, left-side limbs (fore and hind) were shifted inward by *s* (reducing their distance from midline to *h* − *s*), while right-side limbs were shifted outward to *h* + *s*. This modifies the initial stance positions and swing target positions in the body’s coordinate system, altering the geometry of the support polygon formed by the paws. As a result, the distribution of vertical loads and horizontal ground reaction forces becomes biased, shifting the COM’s effective path and inducing curvature through changes in the inverted pendulum force (during two-limb support) and overall torque balance. Lateral shift is treated as a turning asymmetry because it directly breaks left-right symmetry to generate curvature, whereas axial shift is treated separately as a sagittal stride-control mechanism that can be applied across turning strategies to preserve admissible support during turning.

### Axial (sagittal) shift of paw placement as a stride control mechanism

To improve turning stability, we introduced a stride control mechanism implemented as axial (sagittal) shift of paw placement during the swing-to-stance transition. Animals may use such or similar stride adjustment mechanisms during turning, and this adjustment can accompany any of the turning strategies considered here. As illustrated in Fig. 3, the target footfall position is shifted posteriorly relative to the nominal unshifted target in body coordinates. In the baseline model, by contrast, limbs land at fixed anterior target positions relative to the body frame.

**Figure 3:**
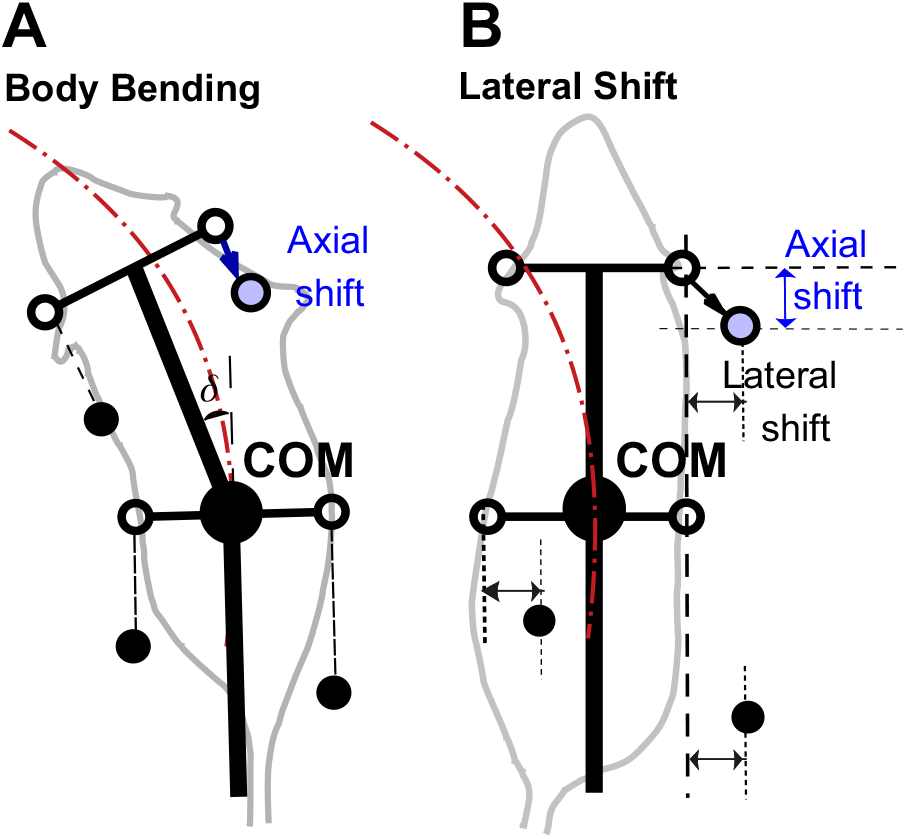
Axial shift implementation in turning strategies. **(A)** Body bending strategy. **(B)** Lateral shift strategy. The Lateral Force case is omitted because this strategy does not alter touchdown geometry or paw-target positions; unlike body bending and lateral shift, its effect is purely dynamic (added perpendicular force) rather than kinematic and is therefore not illustrated by changes in target paw position alone. Black filled circles represent the positions of limbs that are currently in stance phase (ground contact). Open circles indicate default target positions for the limb during the swing phase. Blue circles indicate the adjusted target position in the presence of a non-zero axial shift. The arrows indicate the displacement from the default target (open circle) to the shifted target (blue circle). This posterior shift adjusts the foot placement relative to the body’s center, modifying the support polygon during the turn.

The effect of axial shift is not simply to enlarge or reduce the support polygon. Rather, during turning it reshapes the support geometry and changes which support edge the center-of-mass (COM) trajectory approaches first. Figure 4 illustrates one representative three-limb support phase in which this mechanism can be visualized. In that example, the posterior shift of RF reshapes the support triangle so that the COM trajectory approaches the edge LH-RF earlier, promoting earlier unloading and liftoff of RH. More generally, the same principle applies across turning phases: axial shift alters support geometry and unloading order, while the specific limb and support edge involved depend on the instantaneous support configuration.

**Figure 4:**
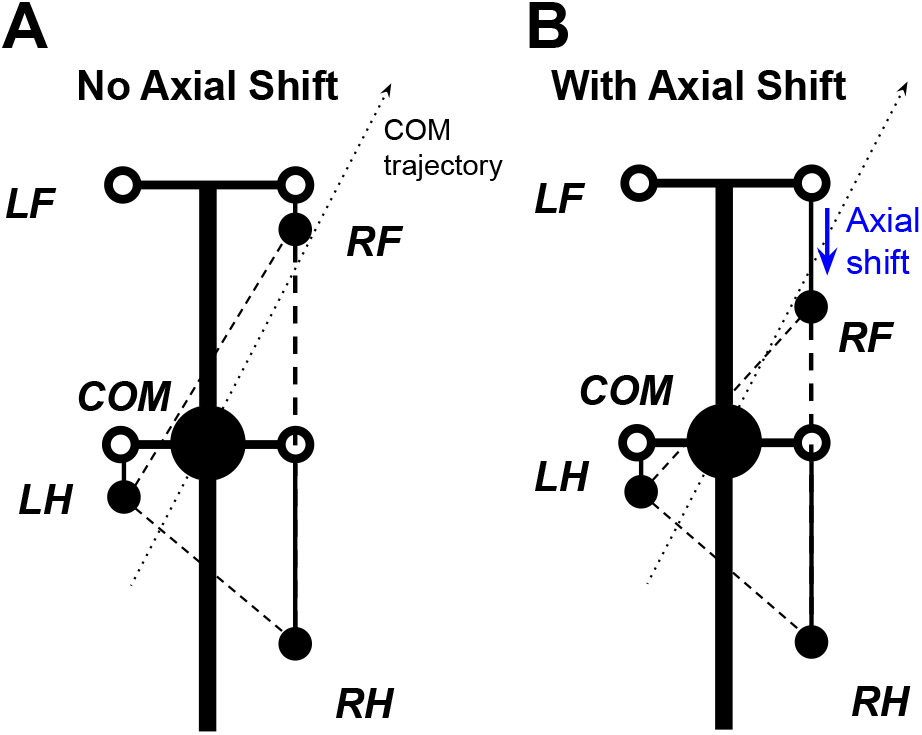
Effect of axial shift on support geometry during turning. Schematic for a representative three-limb support phase with the left forelimb (LF) in swing. **(A)** Without axial shift, the COM trajectory tends to approach the outer support edge RF-RH first, promoting outer-side loss of support. **(B)** With posterior axial shift of the right forelimb (RF), the support triangle is reshaped so that the COM trajectory approaches the edge LH-RF earlier. Because LH-RF is the edge opposite RH, this representative configuration promotes earlier unloading and liftoff of RH, shortening its stance displacement and increasing effective stepping frequency. Thick black lines represent the body axis and simplified body frame. The large black circle denotes the center of mass (COM). Filled black circles indicate actual paw positions in the illustrated phase, while open circles indicate unshifted target paw positions in body coordinates. The dotted arrow indicates the COM trajectory, and the arrow indicates the imposed posterior axial shift of RF.

This earlier unloading shortens the effective stance excursion of the affected limb and advances the next support update, which can increase effective stepping frequency during turning. Repeated across successive steps, this mechanism reduces the tendency of the COM trajectory to progress toward rollover-type loss of support and broadens the range of turning conditions that remain admissible in the model.

In this implementation, the associated adjustment of the forelimb parameter *D*, as described in Methods, is a compensatory modeling choice rather than the primary source of the cadence effect. Because axial shift promotes earlier unloading and liftoff of the affected hindlimb, its stance durations would otherwise become shorter than forelimb stance durations. Reducing the allowable forelimb stance excursion helps preserve fore-hind temporal coordination under axial shift. We therefore attribute the increase in stepping frequency primarily to earlier hindlimb unloading and stance termination caused by the altered support geometry, while the forelimb *D* adjustment serves to maintain coordination.

This interpretation is intended for the low-speed walking regime considered here and should not be taken as a general account of dynamic stability in faster, aerial, or fully three-dimensional turning behaviors.

## Results

Our primary objective was to comparatively investigate the three turning strategies described above, focusing on the maximal achievable turning curvature during stable locomotion and its dependence on velocity. While a certain degree of left-right asymmetry is necessary for turning, each type of asymmetry inherently destabilizes locomotion, imposing specific limitations on achievable curvature and speed.

To address this, we first evaluated the performance of these strategies without stride control. We then compared these results to simulations that included stride control (implemented via an axial shift of the limb‘s footfall target position). As we show, the addition of stride control significantly expands the range of velocities and curvatures available for stable locomotion. Our analysis of curvature as a function of velocity revealed distinct performance profiles for each strategy, delineating regions of instability where the model could not maintain stable locomotion. These regions represent physical constraints inherent to planar walking, where excessive speed or asymmetry can lead to a loss of balance. Similar stability constraints are a primary focus in the design of quadrupedal robotic control systems (Lee et al., 2020).

### Asymmetries destabilize walking without stride control (no axial shift)

To explore the relationship between turning curvature and locomotor velocity for different turning strategies, we constructed corresponding heatmaps for each: body bending (Fig. 5A), lateral force (Fig. 5B), and lateral shift (Fig. 5C). In these maps, successful turns are represented by the shaded areas, whereas the white areas indicate instability and therefore delineate the system’s dynamic boundaries. In the present model, instability refers to the loss of an admissible support or load configuration within the planar walking approximation, not to a fully modeled vertical collapse of the body. A key observation is that none of the turning strategies yield high curvatures when the limbs are set to their default position at the swing-to-stance transition.

**Figure 5:**
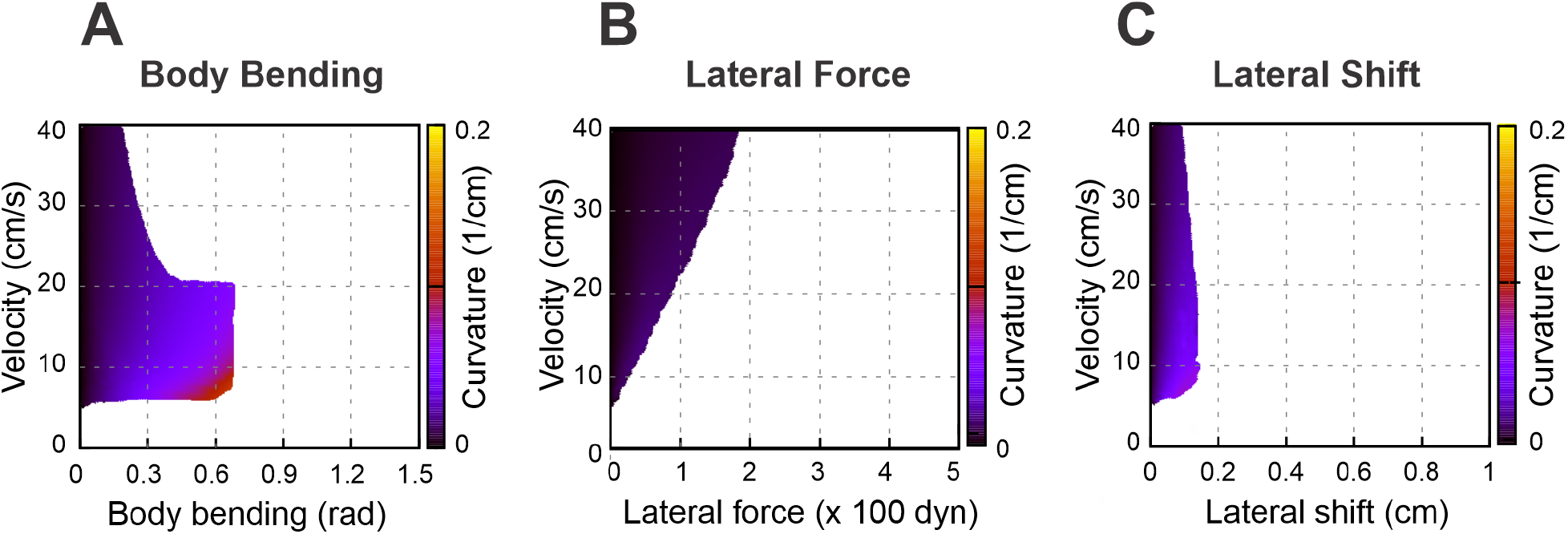
Heatmaps illustrating the turning curvature for three distinct strategies: body bending **(A**), lateral force (**B**), and lateral shift (**C**). These heatmaps show the relationship between turning curvature, velocity (regulated by the propulsion force), and the intensity of each strategy. Shaded areas denote stable walking, while white regions indicate instability, signifying that the model was unable to sustain stable locomotion. For the present mouse-like geometry (*L* = 10 cm, body width 2*h* = 2 cm), 10 cm/s corresponds to 1 body length per second, a lateral shift of 1 cm corresponds to 0.1 body length or 0.5 body widths, and curvature 𝓍 can be interpreted in normalized form as 𝓍*L*; equivalently, turning radius is *R* = 1/𝓍, or *R*/*L* = 1/(𝓍*L*) in body-length units.

White areas in Fig. 5A–C indicate unstable locomotion. When an asymmetry is applied (body bending, an added lateral force, or a lateral limb shift), an unbalanced torque about the body COM (yaw) is produced. That torque rotates the rigid body frame in yaw, so the ground-plane positions of the body-frame anchor points that define propulsive-force directions shift relative to the COM, thereby changing the corresponding moment arms and the stance geometry through which propulsion acts. Because propulsion forces act through those paw positions, rotation tends to reinforce the yaw (a positive feedback loop) which can lead to misalignment of the body axis and the COM movement direction. Stability in our model is geometric: the vertical projection of the COM must remain inside the convex polygon whose vertices are the paws in contact with the ground. As asymmetry grows and the body rotates, the COM projection moves laterally toward the outer side of the turn and the COM projection may cross the support boundary of the outer limbs. Once the COM moves beyond that outer support line the model can no longer produce positive limb loads that keep the body upright – vertical loads go to zero or negative on inner limbs, the remaining outer limbs cannot generate counter-torque, and the body rolls over. A computed negative load is not interpreted as a literal downward pulling ground force. Rather, it indicates that the static balance equations would require such a force to maintain contact, which signals that the corresponding limb can no longer remain in stance and should be treated as unloaded.

### Axial (sagittal) shift at swing-to-stance transition enhances efficiency across all three strategies

To expand the range of stable locomotion velocities and turning capabilities we implemented the stride control via an axial (sagittal) shift—a posterior adjustment of the limb’s footfall target position. While this mechanism generally improved stability, the nature of the enhancement varied distinctly depending on the turning strategy employed. These effects are detailed below and illustrated in Fig. 6 and Fig. 7.

**Figure 6:**
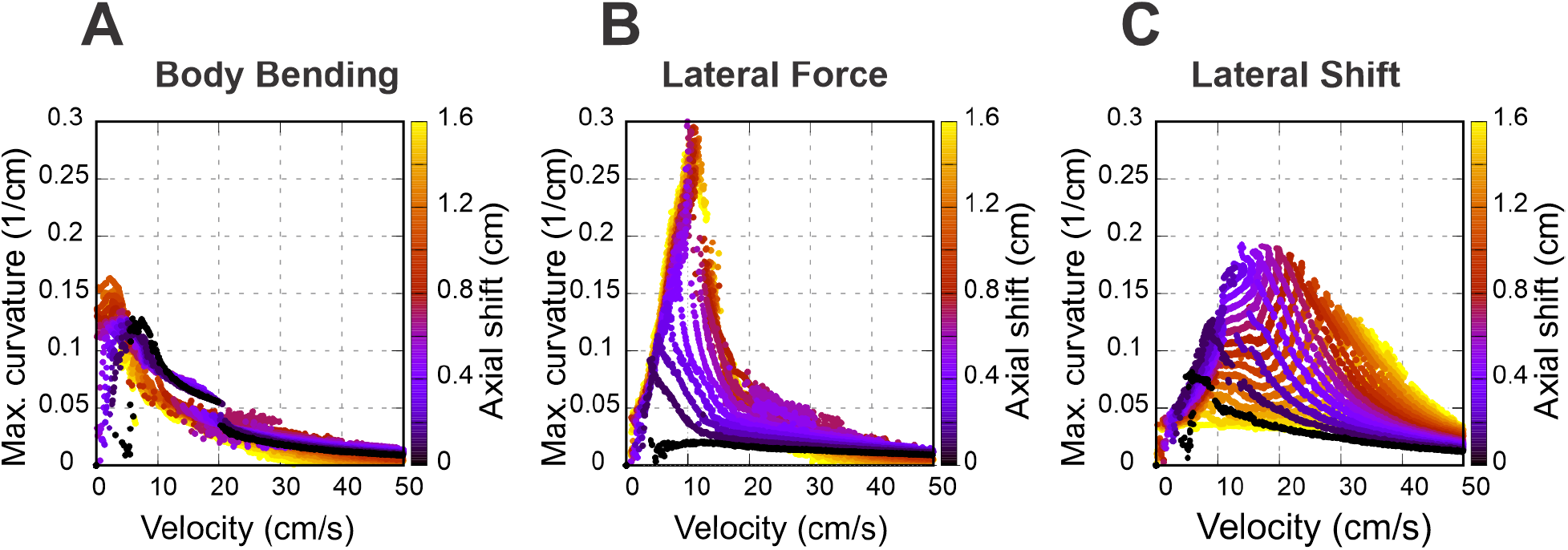
Speed-dependent effects of axial shift on turning performance. Maximum achievable curvature is plotted against velocity for body bending (**A**), lateral force (**B**), and lateral shift (**C**). Color corresponds to the magnitude of the axial shift in centimeters. Note that body bending only allows for sharp turns at low speeds and lateral force achieves the highest overall curvatures at medium speeds; both saturate at ≈1 cm shift, corresponding to 0.1 body lengths (*L*) or 0.5 body widths (2*h*) for the simulated mouse geometry. In contrast, in the lateral shift strategy, the axial shift pushes the performance curve toward higher velocities, peaking at a smaller axial shift of ≈0.4 cm. Compare with the unshifted results in Fig. 5.

**Figure 7:**
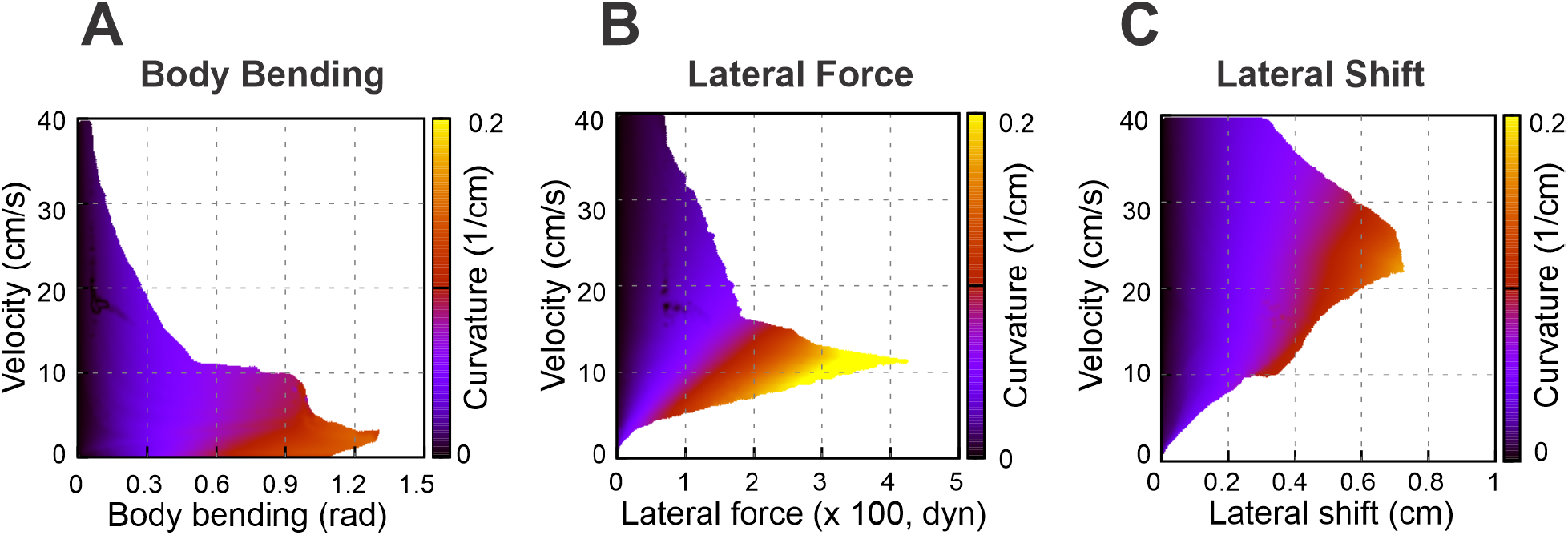
Stability heat maps showing the feasible turning regions in the parameter space of velocity and strategy intensity for selected nonzero axial-shift conditions: 1 cm for body bending (**A**), 1 cm for lateral force (**B**), and 0.4 cm for lateral shift (**C**). These values were chosen as representative nonzero axial-shift conditions to illustrate the substantial stabilizing effect of axial shift seen in Fig. 6. Compare with the zero-shift conditions in Fig. 5 –C.

Specifically, Fig. 6 shows how the maximal turning curvature changes with velocity at different values of axial shift for each turning strategy considered: body bending (Fig. 6A), lateral force (Fig. 6B), and lateral shift (Fig. 6C). Figure 7A-C show the same velocity-strategy heat maps as Fig. 5A-C, but for representative nonzero axial-shift conditions. These axial-shift values were chosen based on the trends in Fig. 6, because they produced substantial changes in the extent and/or position of the stable turning region for each strategy. These can be compared with the corresponding maps in Fig. 5A-C calculated with zero axial shifts.

The most pronounced effect of the axial shift on the body bending strategy (Fig. 7A) is the stabilization of turns at low velocities (<5 cm/s). Without this shift, body bending is ineffective at slow speeds due to instability (Fig. 5A). The axial shift counteracts this, allowing the strategy to function effectively even at very low speeds. The performance benefits for this strategy increase with the magnitude of the shift up to a saturation point of approximately 1 cm, which corresponds to 0.1 body lengths (*L*) or 0.5 body widths (2*h*) for the simulated mouse geometry.

For the lateral force strategy (Fig. 7B), the axial shift facilitates turning primarily at medium speeds (5–5 cm/s). With the axial shift applied, this strategy is capable of generating the highest maximum curvatures of all three methods. Similar to body bending, the efficiency gains for lateral force saturate at an axial shift of approximately 1 cm, corresponding to 0.1 body lengths (*L*) or 0.5 body widths (2*h*) for the simulated mouse geometry, beyond which no further improvement in curvature is observed.

The effect of axial shift on the lateral shift strategy (Fig. 7C) differs qualitatively from the others. Instead of simply extending the range, the axial shift displaces the performance curve, shifting the optimal turning capability from intermediate speeds to higher speeds (15–40 cm/s). Furthermore, the magnitude of axial shift required to peak performance is lower; the turning curvature maximizes at approximately 0.4 cm of axial shift, distinguishing it from the 1 cm saturation point observed in the other two strategies.

The mechanisms underlying these performance limits reflect the interplay between speed and stability. At low velocities, body bending effectively reorients the body axis because static stability is easily maintained. As velocity increases, the centrifugal forces of tight turns disrupt the static support polygon, rendering geometric bending insufficient; here, explicit lateral forces provide the centripetal acceleration needed to actively steer without immediate rollover. At the highest walking speeds, inertial forces dominate, and lateral shift becomes the most viable strategy because it dynamically widens the base of support on the outside of the turn, actively preventing the rollover instability that limits the other two strategies.

The comparative analysis summarized in Fig. 8 demonstrates that no single strategy is universally optimal; instead, the effectiveness of a turning mechanism is fundamentally tied to the animal’s locomotion speed. At low velocities, the body bending strategy leverages static stability to achieve sharp turns through axial flexion. As velocity increases, the dynamic requirements of steering favor the lateral force strategy, which produces the greatest turning sharpness by utilizing forelimb-driven torques. Most notably, the lateral shift strategy becomes the most effective at high speeds, where kinematic adjustments to the base of support are necessary to prevent roll-over instability.

**Figure 8:**
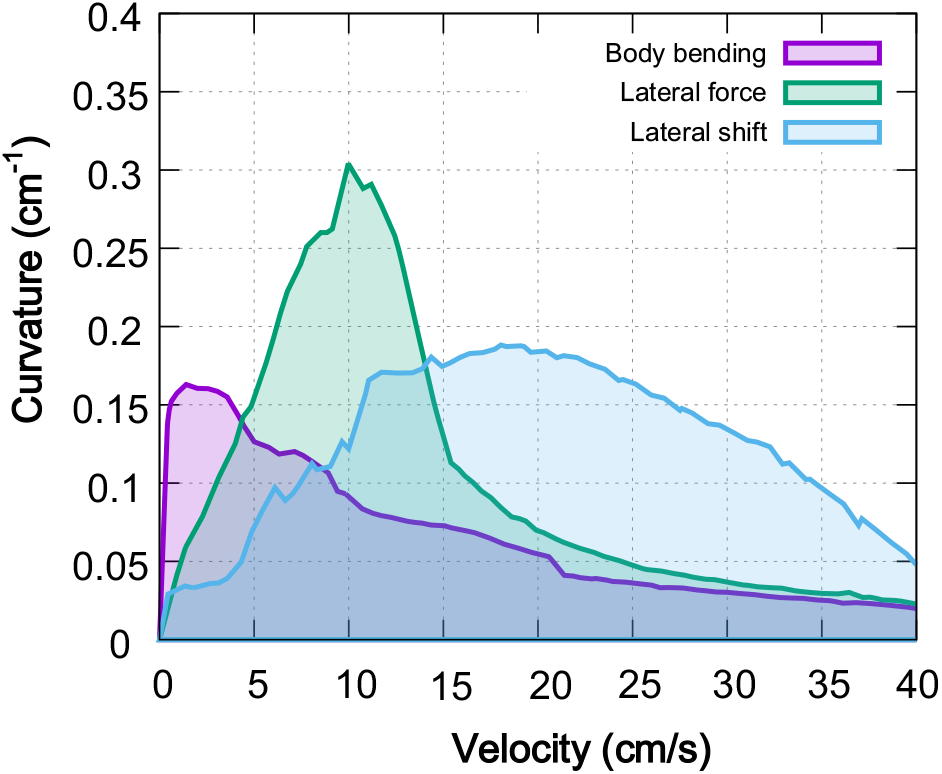
Comparison of maximal turning performance across strategies. Each curve represents the maximum achievable curvature as a function of locomotor velocity for the three primary turning strategies: body bending (purple), lateral force (green), and lateral limb shift (blue). The corresponding shaded regions describe the stable performance envelope under axial shift conditions optimized for each speed. The results reveal a clear speed-dependency: body bending excels at low speeds (<5 cm/s), lateral force generates the highest absolute curvatures at intermediate speeds (5−15 cm/s), and lateral shift maintains superior stability and maneuverability at higher velocities (>15 cm/s).

This suggests that animals may select strategies based on locomotion speed, as seen in robotic models (Wei et al., 2018).

### Combining strategies

In previous sections, we focused on the analysis on three distinct turning strategies. At the same time animals, particularly small quadrupeds like rodents, often combine multiple strategies to perform desired turning under different conditions (Pycock, 1980; Walter, 2003; Gruntman et al., 2007; Usseglio et al., 2020; Haagensen et al., 2022). To investigate whether a combination of different strategies leading to multiple asymmetries could extend the range of achievable curvatures or bridge the performance gaps between individual strategies, we performed brute-force optimization on pairs of strategies. We defined the maximum magnitudes for each mechanism based on the upper limits used in our previous single-strategy heatmaps: 2 rad for body bending, 500 dyn for lateral force, and 1 cm for lateral shift.

For each pair of strategies, we explored their simultaneous activation using a reparameterization of the two-dimensional space of strategy magnitudes. Specifically, we introduced a relative-weight parameter *s* (ranging from 0 to 1) and an overall magnitude parameter *t* (ranging from 0 to 1), such that the first strategy had magnitude *ts Max*_l_ and the second had magnitude *t*(*1* − *s*) *Max*_2_, where *Max*_1_ and *Max*_2_ are the maximal values used for that strategy pair. This parameterization is mathematically equivalent to exploring the full rectangle of pairwise magnitudes defined by those maximal values, while also allowing initialization from the symmetric stable regime at zero asymmetry. We examined 11 discrete relative weightings for each pair (*s* = 0, 0.1,…, 1). For each value of *s*, we swept *t*, propulsion force, and axial shift, and for each speed retained the maximum stable curvature obtained. Thus, each curve in Fig. 9 represents the best performance along one direction in the space of pairwise combinations, and the upper envelope across the curves represents the maximal curvature achieved within the explored space.

**Figure 9:**
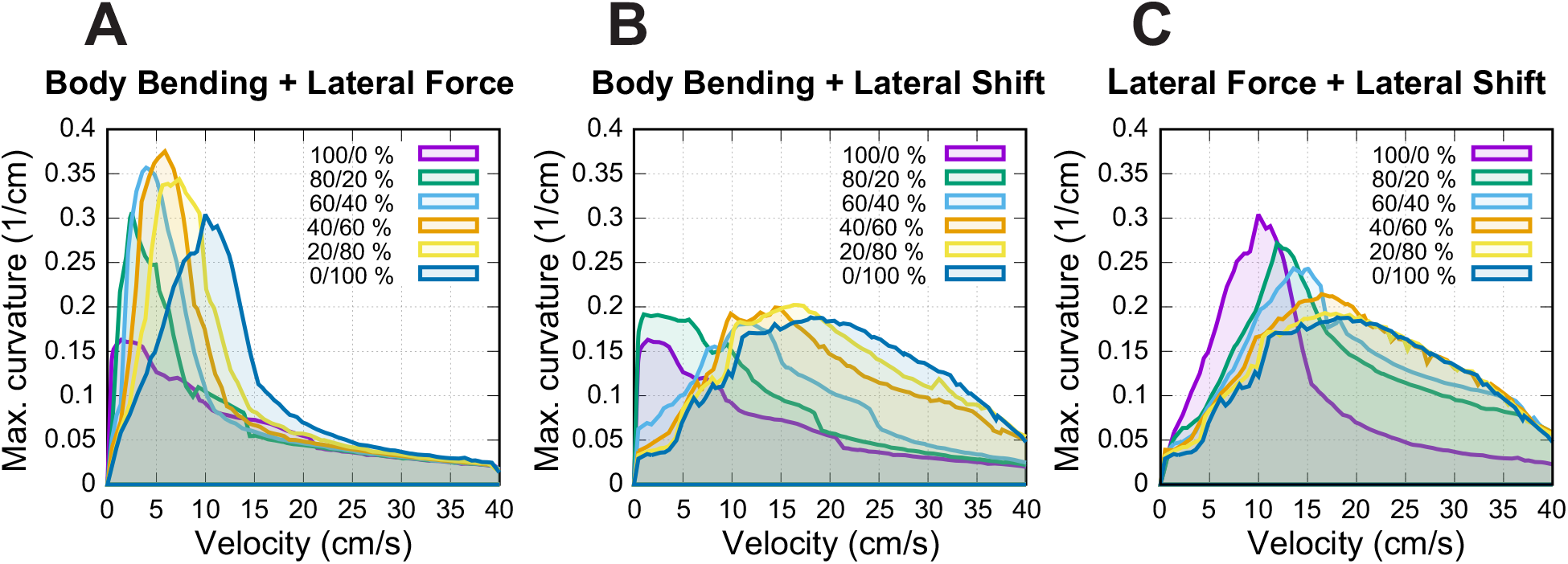
Maximum achievable curvature using combined turning strategies. Each curve represents, for one relative weighting shown in the legend, the maximum stable curvature obtained at each velocity for that pair of simultaneously activated strategies (see text for details). For each relative weighting, the overall combination magnitude, propulsion force, and axial shift were swept to identify the stability boundary. The upper envelope across the curves therefore represents the maximum curvature achieved within the explored space of pairwise combinations.

Our analysis shown in Fig. 9 reveals that combining strategies effectively “fills in the gaps” in performance observed when strategies are used in isolation. While single strategies are dominant in specific speed ranges, their efficacy drops off in the transition zones. The pairwise combinations maintain high curvature in these intermediate zones, and their upper envelope defines a more continuous profile of maximal maneuverability across the velocity spectrum.

The results demonstrate that the combination of multiple strategies via simultaneous activation of different asymmetries allows for sharper turns in transition zones where single-strategy performance typically degrades. This effect is most visible in Fig. 9A, where combining body bending and lateral force produces a result that stands out from the other pairings. In this specific case, the two strategies work together to reach substantially higher turning curvatures than what either method can achieve on its own, pushing the model’s maneuverability to a new peak level at low speed.

In contrast, Fig. 9B shows that combining body bending and lateral shift does not make the turns any sharper (i.e., increase maximal curvature).

Instead, this pairing helps the model produce higher curvatures over a wider range of speeds. Similarly, Fig. 9C shows that the interaction between lateral force and lateral shift mostly serves to bridge the gaps between the two strategies. While this combination helps the model transition smoothly between different modes and fills in performance “valleys”, the upper envelope across the explored pairwise combinations does not exceed the best individual-strategy limits for this pairing.

### Strategy-specific remodeling of limb coordination

The analysis of limb coordination reveals that turning requires a distinct reorganization of gait timing, quantified here by the ratio of duty factors between inner and outer limbs. For the limb-control rules underlying these coordination changes, see the Model Description and Methods sections “Stance-to-swing transition” and “Swing-to-stance transition.” Figure 10 shows that while forelimb asymmetry follows a consistent pattern across all strategies, hindlimb coordination is highly strategy-dependent. In all panels of Fig. 10 the model is performing a left turn, so the inner limbs correspond to the left side and the outer limbs to the right side. The plotted duty-factor ratios therefore directly compare inner-to-outer coordination.

**Figure 10:**
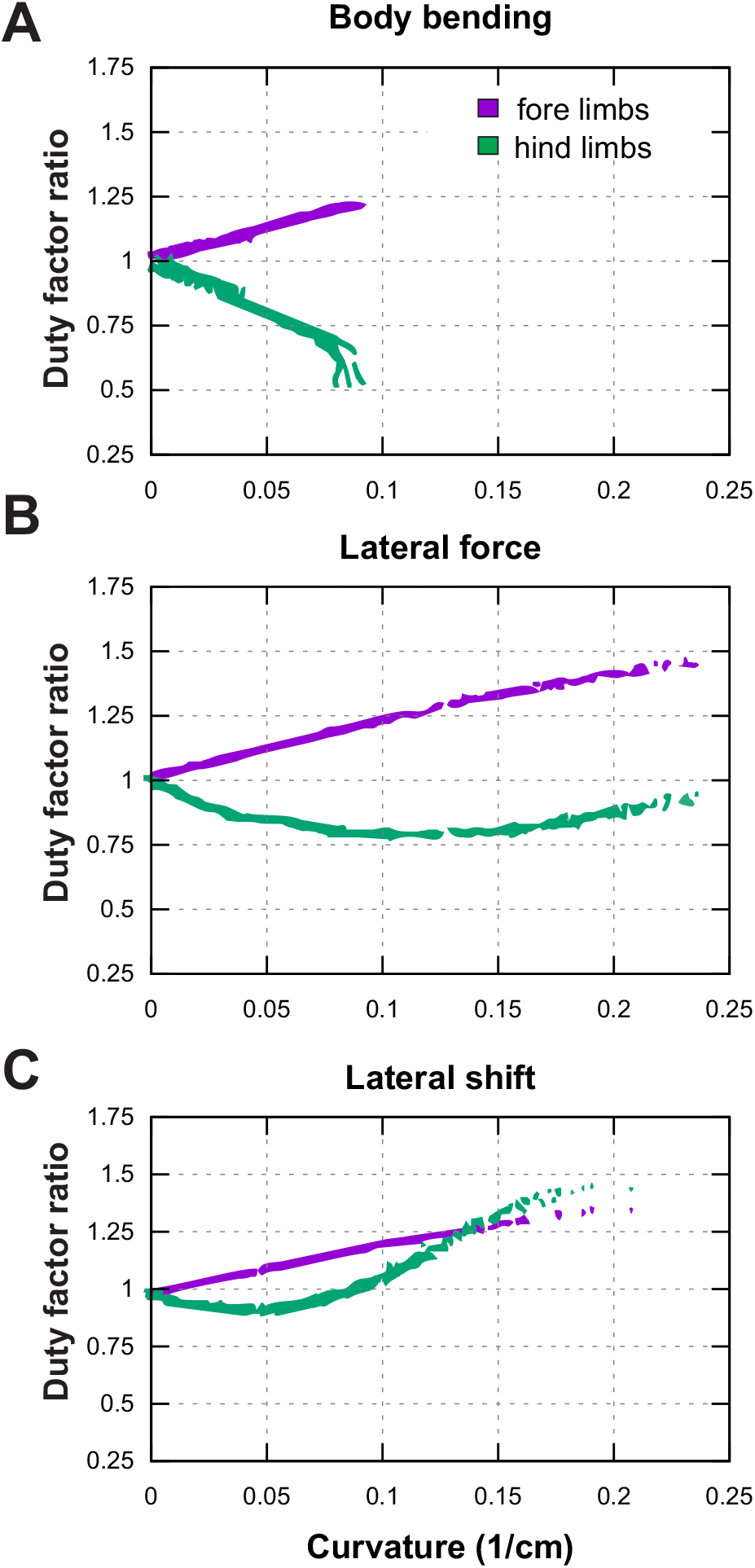
Duty factor ratios (stance duration relative to stride duration) of forelimbs (purple) and hindlimbs (green) as a function of turning curvature. All panels show left-turn examples; therefore the inner limbs are the left limbs and the outer limbs are the right limbs. A ratio > 1 indicates that the inner limb spends a larger fraction of the stride in stance than its outer counterpart (and vice versa for ratios < 1).

Across all three turning strategies—body bending, lateral force, and lateral shift—the forelimbs exhibit a progressive, linear increase in duty factor asymmetry as the turn becomes sharper. This indicates that the inner forelimb consistently spends a longer proportion of the gait cycle in contact with the ground relative to the outer forelimb. In contrast, the hindlimbs display divergent behaviors. In the body bending strategy, hindlimb asymmetry develops in the opposite direction, with the inner limb duty factor decreasing relative to the outer limb as curvature increases. The lateral force strategy shows a non-monotonic pattern: hindlimb asymmetry initially tracks with body bending at low curvatures but reverses trend as the turn tightens. Most notably, the lateral shift strategy produces a sharp, distinct reversal; while it initially resembles the other strategies, at higher curvatures the hindlimb asymmetry spikes dramatically, eventually exceeding the magnitude of asymmetry observed in the forelimbs.

Figure 11 shows that, across the turning conditions illustrated here, the model retains the same basic walking gait organization as in the symmetric baseline case. The principal effect of turning is not a transition to a distinct gait, but a progressive left-right asymmetry in stance and swing timing. In the Body Bending and Lateral Force cases, this asymmetry remains compatible with the baseline walking pattern. In the lateral-shift case at high curvature (Fig. 11D), the most notable additional feature is a substantial overlap between the swing phases of the inner hindlimb (HL) and inner forelimb (FL). This reflects a reduction in diagonality, but is interpreted here as a timing distortion within the same walking gait rather than as a separate gait class.

**Figure 11:**
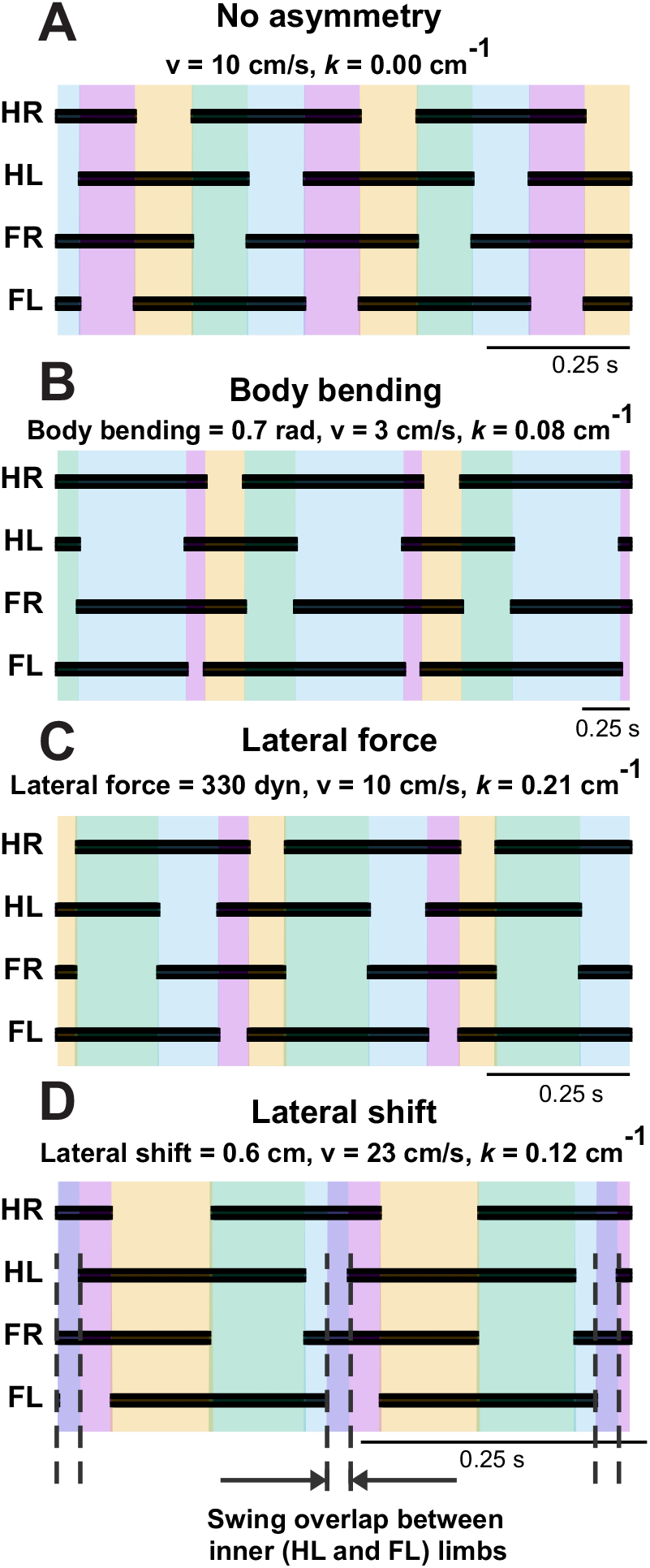
Footfall diagrams for each limb during a left turn: hind right (HR), hind left (HL), fore right (FR), and fore left (FL) limbs during straight walking (**A**) and during turns using body bending (**B**), lateral force (**C**), and lateral shift (**D**), showing temporal sequences of ground contact (black) and swing phases shown by color. In panel D, the highcurvature lateral-shift condition produces a substantial overlap between the swing phases of the inner HL and inner FL, creating a transient interval in which support is borne primarily by the outer limbs. This indicates a marked reduction in diagonality within the same basic walking gait, rather than a transition to a distinct gait class.

Turning was not initiated from a single prescribed touchdown event or gait phase. For each simulation, the model was first allowed to reach a stable periodic straight-walking regime at the selected speed, after which the turning asymmetry was introduced gradually over multiple step cycles.

## Discussion

### Velocity-dependent orchestration of turning strategies

Our results suggest that no single turning strategy is uniformly best across the walking speeds examined here. Instead, the model indicates that the mechanically effective route to curvature depends on locomotor speed: body bending is most effective at low speeds, explicit lateral force performs best at intermediate speeds, and lateral shift becomes comparatively advantageous at higher walking speeds. We interpret this pattern primarily as a speed-dependent interaction among mechanical constraints, support geometry, and stride timing within the model. It is therefore more cautious to view these results as identifying candidate principles for speed-dependent turning control, rather than as direct evidence that animals or the nervous system explicitly switch among discrete strategies in exactly this manner. A useful interpretation of the present framework is that turning acts as an asymmetric modulation of an ongoing locomotor controller, rather than as an entirely separate controller replacing the baseline walking pattern.

At low velocities, the Body Bending strategy dominates by leveraging axial flexibility to reorient the heading direction. This aligns with behavioral studies in rodents showing that spinal curvature is the primary determinant of turning radius during slow exploration (Pycock, 1980). However, as velocity increases, the centrifugal forces associated with sharp turns overwhelm the static stability provided by bending alone. Our model indicates that a transition to the Lateral Force strategy is required. This strategy uses forelimb-driven torque to actively steer the body, effectively “muscling” the system through the trajectory when momentum precludes simple geometric reorientation. Finally, at the highest velocities, the Lateral Shift strategy becomes essential. By widening the base of support on the outside of the turn, this strategy addresses the critical failure mode of high-speed turning: roll-over instability. A possible interpretation is not that the control system simply deprioritizes sharp turning at higher speeds, but that the mechanically admissible range of stable curvatures narrows as speed increases. In this view, the reduction in maximal curvature emerges from an interaction between control and physical constraint: the system may attempt to turn more sharply, but support geometry, load redistribution, and stance timing limit what can be achieved without destabilizing walking.

The biological interpretation of the present results is strongest at the level of turning kinematics and coordination. The model predicts that turning can be supported by left-right locomotor asymmetries, including asymmetric limb timing and body-orientation changes. This is broadly consistent with recent mouse studies showing that directional control is associated with locomotor gait asymmetries and limb-based turning mechanisms (Cregg et al., 2024). By contrast, the more specific assignment of distinct mechanical roles to different limb groups in the present model should be interpreted as a model-based hypothesis rather than a directly established feature of mouse turning behavior. In this context, body bending corresponds to a geometric reorientation of the body axis, whereas the Lateral Force and Lateral Shift mechanisms represent two distinct mechanical routes by which steering-related lateral forces can arise. Because direct turning-force measurements are not analyzed in the present study, the model’s force-redistribution interpretations should be viewed as qualitative hypotheses rather than direct experimental conclusions.

A more direct biological comparison is with turning sharpness rather than speed alone. In Fig. 9, the highest curvatures produced by the model reach approximately 0.3–0.35/cm, corresponding to turning radii of about 3–3.33 cm, or roughly one third of body length for the present mouse-like geometry. This places the model in a regime of very sharp, localized turns. Although direct stride-resolved curvature measurements are scarce in the mouse literature, this scale is consistent with sharp cornering reported experimentally. Zouwen et al. (2021) analyzed freely moving mice making 90° turns in a 40 x 40 cm arena using corner regions of interest of radius 20 cm and a geometrically defined turning point at the corner diagonal. A visual estimate of their trajectories suggests that the sharpest turns also occur with radii on the order of a few centimeters. They further showed that mice transiently brake during such turns, with turn speed scaling with both entry speed and turn angle. Thus, while the experimental measurements are not reported in exactly the same curvature metric used here, they support the conclusion that the sharpest turns produced by the model fall within a biologically plausible range for mouse cornering.

Because inertia, load distribution, and support geometry scale with body mass and body size, the relative advantage of different turning strategies may vary across species; extending the model across mass scales is therefore an important direction for future work.

### Mechanistic role of axial shift: coupling geometry and timing

A critical finding of this study is the stabilizing role of the stride control. In the present model, this control is achieved by shifting the paw backward as the limb transitions from swing to stance. Its stabilizing effect is geometric rather than a simple enlargement or recentring of the support polygon. By reshaping the support geometry, axial shift changes which support edge the COM trajectory approaches first and thereby changes the unloading sequence, helping avoid rollover-type instability during turning. In representative configurations, this also promotes earlier hindlimb unloading and liftoff and thereby alters the timing of support updates.

This interpretation should be understood within the low-speed planar walking regime considered here. Any energetic trade-off or neural regulation of axial shift lies beyond the present model and remains a question for future experimental and computational work.

### Synergy and robustness in multimodal control

The interaction maps (Fig. 9) suggest that the central nervous system likely employs a “mixed-mode” control strategy rather than switching between independent mechanisms. Notably, the simultaneous activation of Body Bending and Lateral Force (Fig. 9A) produces a synergistic effect, yielding maximal turning curvatures that exceed the limits of either strategy in isolation. This additive performance implies that these two mechanisms are mechanically orthogonal and complementary: Body Bending establishes the geometric arc of the turn by reorienting the body axis, while Lateral Force provides the necessary centripetal acceleration to steer the center of mass along that arc. By combining these distinct mechanical advantages, the system achieves a level of maneuverability that neither strategy can support alone.

However, our current computational analysis was limited to the simultaneous implementation of combined strategies. In contrast, biological quadrupeds, including rodents, often utilize sequential implementations of different strategies or modulate speed to leverage specific speed-dependent mechanisms. For instance, animals may initially reduce locomotor speed—or break movement entirely—to execute an abrupt, but stable, turn using body bending before accelerating again (Pycock, 1980; Walter, 2003; Gruntman et al., 2007; Usseglio et al., 2020). Future work will test temporally sequenced steering commands, stride-phase-dependent switching between asymmetries, and state-dependent controller architectures that select turning mechanisms according to speed, curvature demand, and current support configuration.

### Gait remodeling and the “virtual bank”

The divergence in duty factors (Fig. 10) and the emergence of overlapping swing phases (Fig. 11) in the Lateral Shift strategy point to a sophisticated remodeling of the gait cycle. As the turn tightens, the inner limbs must drastically alter their timing to maintain the overall rhythm despite traveling a shorter path length.

Most notably, the synchronization of inner fore- and hindlimb swing phases creates a transient period where the body is supported exclusively by the outer limbs. Mechanistically, this unloading of the inner side allows gravity to accelerate the COM into the turn, functionally mimicking the “banking” seen in motorcycle dynamics or bipedal running (Jindrich et al., 2009). By momentarily yielding support on the inner side, the animal utilizes gravitational torque to counteract centrifugal force. This suggests that the “instability” of overlapping swings is not a breakdown of the gait, but a controlled maneuver to facilitate high-speed turning dynamics. This aligns with the broader biological principle that turning is not merely an asymmetric walking gait, but involves coordinated, descending-driven reorganizations of the locomotor pattern (Usseglio et al., 2020).

### Implications for neural control and robotics

Collectively, these results support the view that quadrupedal turning can be generated through coordinated modulation of trunk posture, limb loading, and foot placement, while the specific neural implementation of these adjustments remains to be determined. These findings are consistent with the general idea that turning may emerge from modulation of an ongoing locomotor controller through changes in trunk posture, limb loading, and foot placement. However, the present model does not identify the neural populations responsible for these adjustments, nor does it distinguish whether different turning behaviors require distinct circuits or different operating regimes of shared circuitry. For that reason, any mapping from the present results to specific spinal or supraspinal populations should be regarded as a hypothesis for future experimental work rather than a conclusion of the current study.

For robotics, this study highlights the limitations of rigid or single-strategy steering controllers. To achieve the agility of biological quadrupeds, robotic platforms must move beyond fixed turning kinematics and implement adaptive, speed-dependent strategy switching. While high-speed balance control is already a recognized challenge in robotics (Hattori et al., 2025), our model underscores that mimicking biological agility requires integrating multiple steering mechanisms that transition dynamically with speed. Specifically, integrating an active spine and variable limb placement policies (analogous to the Lateral Shift) is critical for expanding the stable operating envelope of legged robots in complex, high-speed environments.

## Conclusion

Overall, the model provides a minimal framework for analyzing how geometric, dynamic, and kinematic asymmetries interact with speed to shape quadrupedal turning during walking. The results generate testable hypotheses about the roles of forelimb steering, hindlimb stabilization, and stride-control mechanisms in mouse-like locomotion, while also highlighting opportunities for speed-adaptive steering in quadrupedal robots. At the same time, the conclusions should be interpreted within the limits of a simplified planar walking model.

## Methods

### Overview

Our foundational approach utilized a comprehensive neuromechanical model originally developed by (Molkov et al., 2024). This model served as the core basis for our simulations and subsequent analyses regarding turning strategies. The model’s key components include representations of the central pattern generators (CPGs) for locomotion, basic biomechanics, and a sensory feedback control, allowing for a dynamic and integrated simulation of locomotor and steering behaviors.

The CPG was simplified as a state machine with each Rhythm Generator (RG) existing in one of two possible states at any given time, corresponding to the limb’s stance and swing phases. The body was modeled as a rigid rod (Fig. 1) confined to the horizontal plane at a constant height. This mechanical system consequently possesses three degrees of freedom: two coordinates for the center of mass (COM) in the horizontal plane and the rod’s orientation.

Body movement within our model arises from propulsive forces generated by limbs in the stance phase. We assume these forces are equal in magnitude and aligned with the paw’s displacement during stance in the body’s coordinate system. During the swing phase, a limb does not contribute to propulsion.

Limb loading is defined as the vertical component of the ground reaction force, calculated based on the COM coordinates relative to the positions of the limbs on the ground. For periods with only two limbs on the ground, the body weight cannot be fully supported, so we incorporated the corresponding horizontal components of the inverted pendulum forces into the equations of motion.

Transitions from stance to swing for each limb occur when either the limb’s loading becomes zero or negative, or the limb’s extension surpasses its limit. Swing-to-stance transitions occur based on a feedback control mechanism for swing duration based on potential loss of balance which is defined as inability of the limbs to fully support the body.

A detailed description of each model component follows below.

Parameters in Table 1 were determined based on three criteria. Body dimensions (*L, H, h*), mass (*m*), and maximal limb displacement (*D*) were selected to approximate the geometric scale of a mouse. The gravitational acceleration (*g*) is a standard physical constant. The friction coefficient (*λ*) and nominal propulsion force limits were tuned within the present study to achieve stable baseline walking and enable a broad range of simulated turning maneuvers based on the planar framework established in (Molkov et al., 2024).

**Table 1:**
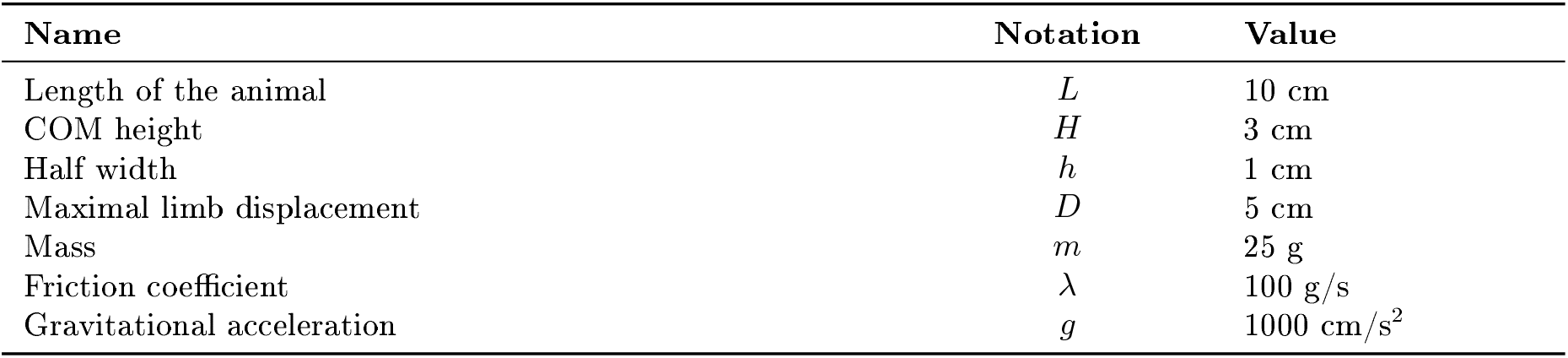
Model parameters. Parameters in this table were determined based on three criteria. Body dimensions (*L, H, h*), mass (*m*), and maximal limb displacement (*D*) were selected to approximate the geometric scale of a mouse. The gravitational acceleration (*g* ) is a standard physical constant. The friction coefficient (*λ*) and nominal propulsion force limits were tuned within the present study to achieve stable baseline walking and enable a broad range of simulated turning maneuvers based on the planar framework established in (Molkov et al., 2024).

| Name | Notation | Value |
| --- | --- | --- |
| Length of the animal | $L$ | 10 cm |
| COM height | $H$ | 3 cm |
| Half width | $h$ | 1 cm |
| Maximal limb displacement | $D$ | 5 cm |
| Mass | $m$ | 25 g |
| Friction coefficient | $\lambda$ | 100 g/s |
| Gravitational acceleration | $g$ | 1000 cm/s<sup>2</sup> |

### Model of the CPG

We implement the CPG as a set of four rhythm generators (RGs), each controlling one limb. Each RG operates in a state machine regime, so that at each moment in time it can be in one of two states: swing or stance. During stance, the end-effector of the limb (paw), which is controlled by the corresponding RG, is assumed to be on the ground hence providing support for the body. During swing, the limbs instantaneously move to their target position relative to the body and await touchdown.

### Model of the body and limbs

For the mouse’s body, we define a rigid frame with length *L* and width 2*h* (Fig. 1). The plane of this frame, also referred to as the *horizontal plane*, is parallel to the ground, and therefore the normal direction to this plane is the *vertical direction*. In terms of mass distribution, the body is considered as a uniform rigid rod of length *L* with the center of mass (COM) at the midpoint of the body. The distance from the horizontal plane to the ground is defined to be the height *H* of the mouse’s COM.

The body weight is supported by the limbs that are in stance. The initial positions of the paws in stance correspond to the ground projections of the front of the body’s frame for the forelimbs and of its center (crossing the COM) for the hind limbs (Fig. 1). The initial stance positions of the left and right limbs are displaced from the body’s centerline by distance *h* to the left and to the right. These initial points serve as target positions in the horizontal plane for the corresponding paws during swing. Here *D* denotes the maximal two-dimensional displacement magnitude of a paw from its touchdown reference position in body coordinates, rather than only the axial component of that displacement. Consequently, a mediolateral offset can cause the displacement limit to be reached earlier even when the axial component of stance excursion is reduced.

### Equations of motion

Let *m* be the mass of the mouse and *g* be the gravitational acceleration. The external forces considered are the gravity force, friction force, and the ground reaction forces. We decompose the forces (where applicable) into components parallel (horizontal) and perpendicular (vertical) to the horizontal plane. Unless otherwise stated, we indicate ***vectors by bold italic letters*** and their *magnitudes* using the same notations in an *italic font*. In the following subsections, we decompose the ground reaction forces into their components in the horizontal plane (as those relevant for planar movements of the body) and their vertical components as those providing body weight support.

Let ***F*** _*i*_ denote the component of ground reaction forces in the horizontal plane (2D vector) for limb *i* (1 for the left fore (LF), 2 for the right fore (RF), 3 for the left hind (LH), and 4 for the right hind (RH) limbs). Specifically, we assume that every paw touching the ground creates a propulsion force ***F*** _*i*_ with magnitude *F*_0_ (same for all limbs on the ground) directed along the paw’s displacement from its initial position during stance in the body’s coordinate system (see Fig. 12B for an illustration). *F*_0_ is the main parameter of the model affecting the locomotor speed. Hereinafter, we refer to this parameter as *propulsion force* and use it as one of the control parameters in all our simulations.

**Figure 12:**
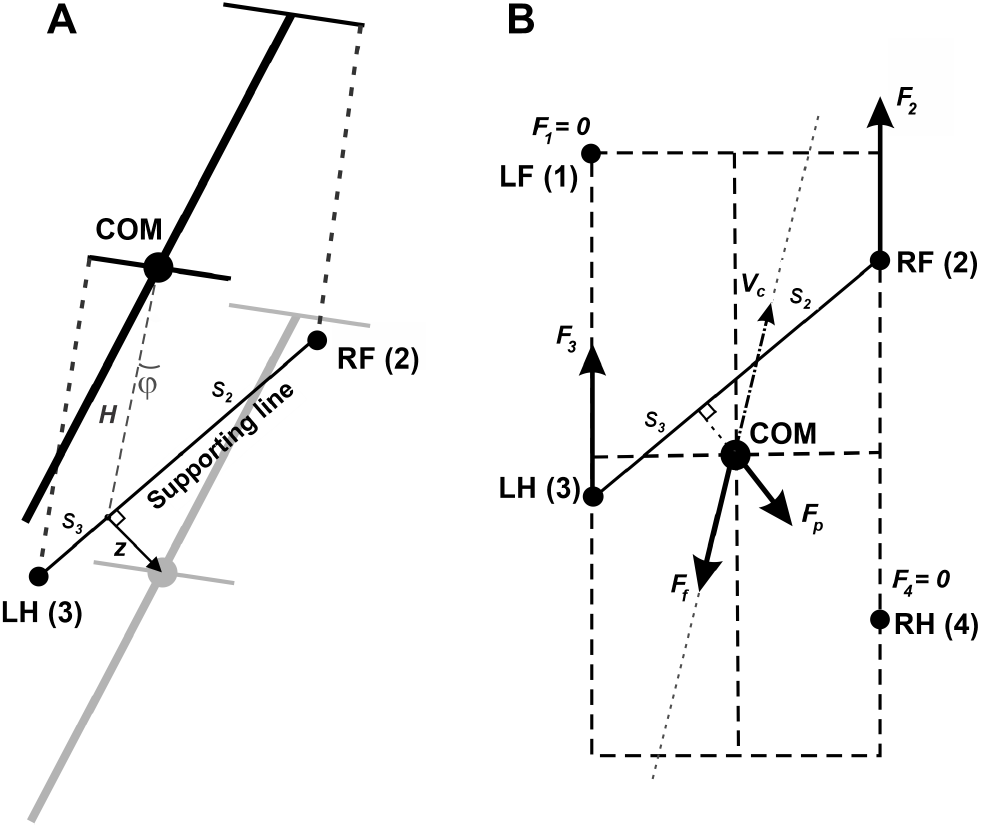
**A.**Inverted pendulum dynamics during two-leg support. During two-leg support phases, the body weight cannot be fully supported by the limbs on the ground. In this case, we describe the effect of the gravitational force on the horizontal center of mass (COM) movement by the linearized inverted pendulum model. In the example shown, the right fore-(RF (2)) and left hind-(LH (3)) limbs are on the ground providing support, while the left fore-and right hindlimbs (not shown) are in swing. Since the COM is displaced from the supporting line (***z*** is the COM displacement vector in the horizontal plane, *ϕ* is the corresponding angle between the projection of the COM onto the line of support and the vertical), the gravitational force creates a rolling torque about this line. We approximate this torque as an equivalent horizontal force pushing the COM in the direction perpendicular to the supporting line (along ***z***). See text for details. **B**. A force diagram in the horizontal plane for the same body configuration as shown in panel A. LF and RH limbs are in swing and do not create propulsive forces. LH and RF push forward with forces ***F***_2_ and ***F***_3_ equal in magnitude and directed along the body axis. The kinetic friction force ***F***_*f*_ is applied at the COM and directed against the COM velocity ***v***_*c*_. The inverted pendulum force ***F***_*p*_ is orthogonal to the line of support connecting LH and RF and pulling the COM away from the line of support. ***F***_*p*_ = **0** in case there are more than two limbs on the ground. This sketch is for illustrative purposes only. While it depicts the general concept, it does not reflect the exact proportions or relative force magnitudes of a real mouse. Reproduced from Molkov et al. (2024).

During locomotion, energy losses are associated with various factors many of which are not explicitly represented in our simplified model. To account for energy dissipation, we introduce an equivalent viscous friction force that linearly depends on the velocity, i.e., ***F*** _*f*_ = *λ****v***_*c*_, where ***v***_*c*_ is the COM velocity vector in the horizontal plane and *λ* is the coefficient of kinetic friction (see Table 1 for parameter values).

When only two limbs support the body, as shown in Fig. 12A, the full static support conditions used for three-or four-limb support cannot in general be satisfied. We therefore use a planar effective-force approximation. The two grounded paws define a support line segment that acts as the pivot line of an inverted-pendulum approximation. Let ***z*** be the distance vector from this support line to the COM projection on the ground (Fig. 12A). The corresponding linearized dynamics are *d*^2^ ***z***/*dt* ^2^ = *g****z***/*H*. Multiplying this acceleration by the mass *m* yields the effective “inverted-pendulum” force ***F*** _*p*_ = *mg* ***z***/*H*, which acts perpendicular to the support line and represents the destabilizing effect of gravity within this reduced planar description. During stance, grounded paws remain fixed in ground coordinates until liftoff, while the body moves relative to those fixed contacts. If more than two limbs are on the ground, ***F***_*p*_ =**0**.

By Second Newton’s Law, the velocity ***v***_*c*_ obeys the following equation (Fig. 12B):

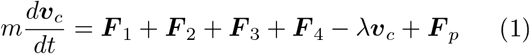

Since the propulsion forces are displaced from the axis of the body, they will contribute to the yaw torque leading to the rotation of the body around the COM in a horizontal plane. The moment of inertia *I* of a uniform rod of length *L* about the COM is *I* = *mL*^2^ /12.

The gravity force is applied at the COM, thus creating zero torque. Let ***r***_*c*_ be the position vector of the COM in the horizontal plane, and ***r***_*i*_ be the position vector of paw *i* on the ground. Then, the yaw torque that each propulsion force creates is ***M*** _*i*_ = ***F*** _*i*_ *×* (***r***_*i*_ – ***r***_*c*_). The friction torque *M* _*f*_ of the rod can be calculated as *M* _*f*_ = *λIω*/*m*, where *ω* is the angular velocity of the body rotation about the COM. By Second Newton’s Law in angular form, the following differential equation describes the rotation of the body about the COM in the horizontal plane:

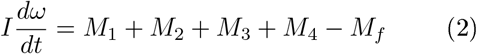

### Weight bearing forces

To calculate the vertical components of the ground reaction forces (***G***_*i*_ ) in each limb that we also refer to as weight-bearing or supporting forces, we consider different possible situations depending on the number of limbs in stance.

#### Support by more than two limbs

We assume that the body frame always remains in the horizontal plane and, therefore, is not pitching or rolling. Therefore, the total torque created by the supporting forces should be balanced, i.e.

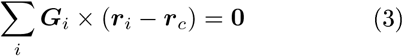

In addition, for the body to remain in the horizontal plane (not move in the vertical direction), the total supporting force should be equal in magnitude to the gravitational force:

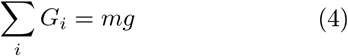

This equality holds in the multi-limb support calculations because body height *H* is fixed and the body has no vertical acceleration.

Finally, ground reaction forces are zero for all limbs in swing.

Note that Eq. (3) contains two equations, because the vector products in the sum have two non-trivial components. Thus, Eqs. (3) and (4) define a linear system of three equations for the supporting forces. This system has a unique solution in the case of three-leg support (provided that the three paws are not positioned on a single straight line). In the case of four-leg support an additional constraint is necessary, as different weight distributions that satisfy Eqs. (3) and (4) are possible. For four-leg support, we choose the solution that distributes load as evenly as possible. This is a regularizing assumption introduced only to obtain a unique solution and should not be interpreted as a claim that biological animals necessarily equalize load in the same way. Particularly, we find a solution of Eqs. (3) and (4) that minimizes Σ_*i*_ ***G***_*i*_ ^2^ using the method of Lagrange multipliers.

For a limb to stay on the ground, the upward component of its ground reaction force must be positive. Geometrically, the system of Eqs. (3) and (4) has positive solutions only if the COM is located inside the triangle (or quadrangle) formed by the limbs in contact with the ground. Whenever the COM falls on the edges of this triangle (quadrangle), the supporting force (load) in the limb opposite to that edge reduces to zero, indicating that the limb needs to be lifted.

#### Support by two limbs

In case of only two limbs being on the ground, the system of Eqs. (3) and (4) does not generally have solutions, meaning that the body weight cannot be fully supported (unless the COM is precisely above the line of support, see Fig. 12). In this two-limb-support approximation, total vertical load is therefore not constrained to remain equal to body weight and can decrease as the COM moves away from the equilibrium configuration relative to the support line. Here we take the case of the diagonal limbs in stance as an example. Consider the left hind-(i = 3) and right fore-(i = 2) limbs are supporting and the other two limbs (i = 1 and 4) are in swing (as shown in Fig. 12), that is, ***G***_l_ = ***G*** _**4**_ = 0. The movement of this frame in the direction perpendicular to the supporting line will follow the inverted pendulum dynamics as illustrated in Fig. 12. Let *v*_*p*_ be the COM velocity component in the direction perpendicular to the supporting line. By taking the centripetal force into account, for the vertical component of the ground reaction force we have:

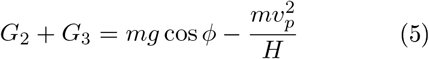

To find the load distribution over limbs 2 and 3 we assume that there is no pitch in the direction of the supporting line, so the torque created by ***G*** _**2**_ and ***G*** _**3**_ about the projection of the COM onto the line of support must be zero. Specifically,

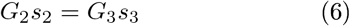

where *s*_2_ and *s*_3_ are the segments of the line of support between the projection of the COM on it and the positions of paws 2 and 3, respectively (see Fig. 12). After solving the linear system defined by Eqs. (5) and (6), we find ***G*** _**2**_ and ***G*** _**3**_ for the case that the body is supported by limbs 2 and 3 only. Similarly, we find the vertical components of the ground reaction forces (also referred to as weight-bearing forces and limb load) for any other pair of supporting limbs during 2-leg support phases.

### Stance-to-swing transition

The transition from stance to swing (lift off) for each limb is governed by local proprioceptive feedback. Modeled after mechanisms observed in biological quadrupedal locomotion (Ekeberg et al., 2005; Molkov et al., 2024), the rhythm generator for a specific limb switches from the stance state to the swing state if either of the following two conditions is met:

#### Limb Unloading (Force-dependent feedback)

The limb is lifted if its load—defined as the vertical component of the ground reaction force—drops to zero or becomes negative (***G*** _*i*_< 0). This rule ensures that a limb does not lift while it is essential for supporting the body’s weight, mimicking the inhibitory feedback from ankle extensor load receptors (e.g., Golgi tendon organs) that prevents swing initiation during substantial weight bearing.

#### Limb Extension (Geometry-dependent feedback)

The limb is lifted if it reaches its maximum posterior displacement limit. The model monitors the displacement of the paw ( |Δ***r***_*i*_ |) relative to its initial touchdown position in the body’s coordinate system. If this displacement exceeds a predefined maximum limb displacement (*D*), the limb is forced to lift ( |Δ***r***_*i*_ | > *D*). This mimics the signal from hip flexor stretch receptors which triggers the flexor burst to initiate swing when the limb is fully extended (Ekeberg et al., 2005).

### Swing-to-stance transition

The transition from swing to stance (touchdown) is governed by the need to maintain static stability. This phase transition is triggered by a “loss of balance” mechanism, which ensures that a swinging limb is grounded immediately if the body’s stability is threatened.

In this model, loss of balance is defined as a negative rate of change of the total load (total vertical ground reaction force, *G*_tot_ = Σ_*i*_ **G**_*i*_). When the total load begins to decrease (i.e., ***dG*** _*tot*_ /***dt*** < 0), indicating that the current support polygon is insufficient to maintain the body’s height, the swing phase must be terminated.

If multiple limbs are currently in the swing phase when this condition is met, the system selects the limb that has been swinging for the longest time to transition to stance. This selection criterion prioritizes the limb that is furthest along in its swing trajectory, ensuring a more natural and efficient recovery of stability.

### Axial (sagittal) shift

To enable precise control over stride length and stepping frequency during locomotion, we incorporated an axial shift mechanism into the model. This strategy involves a controlled rearward displacement of the target paw position along the body’s longitudinal axis at the transition from swing to stance. Specifically, upon initiating stance, the initial paw placement is shifted backward by a variable amount (a control parameter we refer to as *axial shift*) relative to its default unperturbed position in the body coordinate system. This axial shift effectively shortens the stride length without altering the propulsion force magnitude, thereby allowing modulation of gait parameters such as effective step frequency and overall speed.

In implementing axial shift, we also reduce the fore-limb displacement limit *D* by the same amount as the imposed posterior shift. This is a compensatory modeling choice introduced to preserve fore-hind temporal coordination under axial shift, not the primary source of the increase in effective stepping frequency. In the present model, that cadence effect is attributed primarily to earlier hindlimb unloading and stance termination caused by the altered support geometry, whereas the forelimb *D* adjustment serves only to maintain coordination.

### lmplementation of asymmetries for turning

To investigate turning behaviors, we extended the baseline symmetrical model by implementing three distinct biomechanical asymmetries. These strategies introduce controlled deviations from bilateral symmetry to generate the rotational moments required for changing direction.

### Body Bending (geometric asymmetry)

This strategy mimics the spinal flexion observed in biological quadrupeds during turning. In the baseline model, the body is a single rigid rod. To implement bending, we introduce an angular deviation, *δ* (ranging from 0 to 1 radians), between the front and rear segments. The body is effectively treated as two segments where the front segment is rotated by *δ*/2 counter-clockwise (clockwise) and the rear segment is rotated by *δ*/2 clockwise (counter-clockwise) for a left (right) turn. The *δ*/2 construction is a symmetric geometric parameterization around the COM chosen to represent bending within a COM-centered rigid-body framework; it is not intended as a literal anatomical claim that separate trunk segments in the animal independently rotate by exactly *δ*/2. This geometric transformation reorients the limb attachment points in the body coordinate system. Consequently, the swing targets and the resulting support polygon are shifted, generating a yaw torque through the redistribution of ground reaction forces without altering the magnitude of propulsion.

### Lateral Force (dynamic asymmetry)

This strategy simulates the active generation of lateral ground reaction forces, primarily used for steering. We applied a lateral force vector perpendicular to the body axis, directed inward toward the center of the turn (e.g., leftward for a left turn). This force is applied exclusively to the forelimbs to mimic their steering role, while the hindlimbs receive zero lateral force component. The addition of this perpendicular component modifies the horizontal equations of motion by introducing a centripetal force in Eq. (1) and an asymmetric yaw torque in Eq. (2), thereby driving trajectory curving and rotation of the body.

### Lateral Shift (kinematic asymmetry)

This kinematic asymmetry involves offsetting the mediolateral placement of the limbs relative to the body’s midline specifically at the moment of the swing-to-stance transition. In the baseline model, limbs are displaced by a fixed half-width *h* from the center (see Table 1). For a turn, we introduce a lateral offset parameter *s* that modifies the target touchdown coordinates. Specifically, inner limbs (e.g., left) are shifted inward, reducing their distance from the midline to *h* − *s*, while outer limbs (e.g., right) are shifted outward, increasing their distance from the midline to *h* + *s*. This shift alters the geometry of the support polygon and the direction of the propulsion forces. Because propulsion is directed along the paw displacement vector, shifting the starting position naturally introduces a lateral component to the propulsion force vector during stance.

### Calculation of turning curvature

To quantify turning performance, we calculated the signed curvature of the locomotor trajectory using Menger curvature. Menger curvature is a geometric measure defined for a set of three distinct points as the reciprocal of the radius of the unique circumcircle passing through them (ϰ = 1/*R*). This approach allows for a precise, discrete approximation of curvature without requiring a continuous path derivative.

Turning curvature was quantified using Menger curvature computed from the center-of-mass positions at three consecutive touchdowns of the left forelimb. This yields a stride-by-stride estimate of the signed turning radius during locomotion.

For each simulation, the reported value corresponds to the stabilized curvature measured after transient behavior had decayed. Positive values of ϰ indicate left turns, negative values indicate right turns, and larger magnitudes correspond to tighter turns (1/ ϰ).

## Data and Code Availability

The computational model, simulation results, and interactive visualizations are available on the publication companion site at https://math.gsu.edu/ymolkov/turning.

## Acknowledgements

This work was supported by NSF CRCNS grant # 2113069 and NIH/NINDS grants R01NS130799 and R01NS110550 to IAR.

## References

Bertram, J. E. A., ed. (2016). Understanding mammalian locomotion: concepts and applications. Hoboken, New Jersey: Wiley-Blackwell. 1 p. DOI: 10.1002/9781119113713.

Cregg, J. M., R. Leiras, A. Montalant, P. Wanken, I. R. Wickersham & O. Kiehn (2020). “Brainstem neurons that command mammalian locomotor asymmetries”. In: Nature Neuroscience 23.6, pp. 730–740. DOI: 10.1038/s41593-020-0633-7.

Cregg, J. M., S. K. Sidhu, R. Leiras & O. Kiehn (2024). “Basal ganglia–spinal cord pathway that commands locomotor gait asymmetries in mice”. In: Nature Neuroscience 27.4, pp. 716–727. DOI: 10.1038/s41593-024-01569-8.

Danner, S. M., N. A. Shevtsova, A. Frigon & I. A. Rybak (2017). “Computational modeling of spinal circuits controlling limb coordination and gaits in quadrupeds”. In: eLife 6, e31050. DOI: 10.7554/eLife.31050.

Danner, S. M., S. D. Wilshin, N. A. Shevtsova & I. A. Rybak (2016). “Central control of interlimb coordination and speed-dependent gait expression in quadrupeds”. In: The Journal of Physiology 594.23, pp. 6947–6967. DOI: 10.1113/JP272787.

Ekeberg, Ö. & K. G. Pearson (2005). “Computer simulation of stepping in the hind legs of the cat: an examination of mechanisms regulating the stance-to-swing transition”. In: Journal of Neurophysiology 94.6, pp. 4256–4268. DOI: 10.1152/jn.00065.2005.

Frigon, A., T. Akay & B. I. Prilutsky (2021). “Control of Mammalian Locomotion by Somatosensory Feedback”. In: Comprehensive Physiology 12.1, pp. 2877–2947. DOI: 10.1002/cphy.c210020.

Full, R. J. & D. E. Koditschek (1999). “Templates and anchors: neuromechanical hypotheses of legged locomotion on land”. In: The Journal of Experimental Biology 202 (Pt 23), pp. 3325–3332. DOI: 10.1242/jeb.202.23.3325.

Grillner, S. (2006). “Biological pattern generation: the cellular and computational logic of networks in motion”. In: Neuron 52.5, pp. 751–766. DOI: 10.1016/j.neuron.2006.11.008.

Gruntman, E., Y. Benjamini & I. Golani (2007). “Coordination of steering in a free-trotting quadruped”. In: Journal of Comparative Physiology A 193.3, pp. 331–345. DOI: 10.1007/s00359-006-0187-5.

Haagensen, T., J. L. Gaschk, J. T. Schultz & C. J. Clemente (2022). “Exploring the limits to turning performance with size and shape variation in dogs”. In: Journal of Experimental Biology 225.21, jeb244435. DOI: 10.1242/jeb.244435.

Hattori, S., S. Suzuki, A. Fukuhara, T. Kano & A. Ishiguro (2025). “Bicycle-inspired simple balance control method for quadruped robots in highspeed running”. In: Frontiers in Robotics and AI 11, p. 1473628. DOI: 10.3389/frobt.2024.1473628.

Holmes, P., R. J. Full, D. Koditschek & J. Guckenheimer (2006). “The Dynamics of Legged Locomotion: Models, Analyses, and Challenges”. In: SIAM Review 48.2, pp. 207–304. DOI: 10.1137/S0036144504445133.

Jindrich, D. L. & M. Qiao (2009). “Maneuvers during legged locomotion”. In: Chaos (Woodbury, N.Y.) 19.2, p. 026105. DOI: 10.1063/1.3143031.

Lee, J., J. Hwangbo, L. Wellhausen, V. Koltun & M. Hutter (2020). “Learning quadrupedal locomotion over challenging terrain”. In: Science Robotics 5.47, eabc5986. DOI: 10.1126/scirobotics.abc5986.

McCrea, D. A. & I. A. Rybak (2008). “Organization of mammalian locomotor rhythm and pattern generation”. In: Brain Research Reviews 57.1, pp. 134–146. DOI: 10.1016/j.brainresrev.2007.08.006.

Molkov, Y. I., G. Yu, J. Ausborn, J. Bouvier, S. M. Danner & I. A. Rybak (2024). “Sensory feedback and central neuronal interactions in mouse locomotion”. In: Royal Society Open Science 11.8, p. 240207. DOI: 10.1098/rsos.240207.

Müller, V. C. & M. Hoffmann (2017). “What Is Morphological Computation? On How the Body Contributes to Cognition and Control”. In: Artificial Life 23.1, pp. 1–24. DOI: 10.1162/ARTL_a_00219.

Orlovskĭ, G. N. (1999). Neuronal control of locomotion: from mollusc to man. In collab. with T. G. Deliagina & S. Grillner. Oxford: Oxford University Press. 1 p. DOI: 10.1093/acprof:oso/9780198524052.001.0001.

Owaki, D., T. Kano, K. Nagasawa, A. Tero & A. Ishiguro (2013). “Simple robot suggests physical interlimb communication is essential for quadruped walking”. In: Journal of The Royal Society Interface 10.78, p. 20120669. DOI: 10.1098/rsif.2012.0669.

Pfeifer, R., J. C. Bongard & R. Brooks (2007). How the body shapes the way we think: a new view of intelligence. In collab. with S. Grand. A Bradford book. Cambridge, Massachusetts London: The MIT Press. 394 pp.

Pycock, C. (1980). “Turning behaviour in animals”. In: Neuroscience 5.3, pp. 461–514. DOI: 10.1016/0306-4522(80)90048-2.

Rybak, I. A., N. A. Shevtsova, S. N. S. N. Markin, B. I. Prilutsky & A. Frigon (2024). “Operation regimes of spinal circuits controlling locomotion and the role of supraspinal drives and sensory feedback”. In: eLife 13, RP98841. DOI: 10.7554/eLife.98841.

Rybak, I. A., K. J. Dougherty & N. A. Shevtsova (2015). “Organization of the Mammalian Locomotor CPG: Review of Computational Model and Circuit Architectures Based on Genetically Identified Spinal Interneurons”. In: eNeuro 2.5, ENEURO.0069–15.2015. DOI: 10.1523/ENEURO.0069-15.2015.

Rybak, I. A., N. A. Shevtsova, J. Audet, S. Yassine, S. N. Markin, B. I. Prilutsky & A. Frigon (2025). “Operation of spinal sensorimotor circuits controlling phase durations during tied-belt and split-belt locomotion after a lateral thoracic hemisection”. In: eLife 13, RP103504. DOI: 10.7554/eLife.103504.3.

Stuart, D. G. & H. Hultborn (2008). “Thomas Graham Brown (1882–1965), Anders Lundberg (1920–), and the neural control of stepping”. In: Brain Research Reviews 59.1, pp. 74–95. DOI: 10.1016/j.brainresrev.2008.06.001.

Usseglio, G., E. Gatier, A. Heuzé, C. Hérent & J. Bouvier (2020). “Control of Orienting Movements and Locomotion by Projection-Defined Subsets of Brainstem V2a Neurons”. In: Current Biology 30.23, 4665–4681.e6. DOI: 10.1016/j.cub.2020.09.014.

° Walter, R. M. (2003). “Kinematics of 90° running turns in wild mice”. In: Journal of Experimental Biology 206.10, pp. 1739–1749. DOI: 10.1242/jeb.00349.

Wei, Z., G. Song, H. Sun, Q. Qi, Y. Gao & G. Qiao (2018). “Turning strategies for the bounding quadruped robot with an active spine”. In: Industrial Robot: An International Journal 45.5, pp. 657–668. DOI: 10.1108/IR-06-2018-0119.

Zouwen, C. I. van der, J. Boutin, M. Fougère, A. Flaive, M. Vivancos, A. Santuz, T. Akay, P. Sarret & D. Ryczko (2021). “Freely Behaving Mice Can Brake and Turn During Optogenetic Stimulation of the Mesencephalic Locomotor Region”. In: Frontiers in Neural Circuits 15, p. 639900. DOI: 10.3389/fncir.2021.639900.

